# Non-neutralizing antibody responses to vesicular stomatitis virus-vectored influenza A virus vaccines correlate with protection

**DOI:** 10.1101/2022.11.02.514911

**Authors:** Kevin Wittwer, Gert Zimmer, Marcel Rommel, Yvonne Krebs, Svetlana Rezinciuc, Bevan Sawatsky, Christian K. Pfaller

**Affiliations:** Research Group for Pathogenesis of Respiratory Viruses, Division of Veterinary Medicine, Paul-Ehrlich-Institute, 63225 Langen, Germany; Institute of Virology and Immunology, 3147 Mittelhäusern, Switzerland; Research Group for Gene Modification in Stem Cells, Division of Veterinary Medicine, Paul-Ehrlich-Institute, 63225 Langen, Germany; Department of Molecular Medicine, Mayo Clinic, Rochester, MN 55905, United States

**Keywords:** influenza A virus, VSV replicon vaccine, humoral immunity, IgG subclasses, correlates of protection, subtype-independent, Fc-gamma receptor, ADCC, ADCP

## Abstract

Seasonal influenza virus infections remain a major global public health burden. In addition, influenza A virus (IAV) exhibits high pandemic potential through zoonotic spread from avian hosts to humans. Currently licensed IAV vaccines are mainly directed against the immuno-dominant surface protein hemagglutinin (HA). Since HA is antigenically highly variable, IAV can escape vaccine-derived immunity through antigenic drift. However, vaccine preparations such as the live-attenuated influenza vaccine (LAIV) also contain other more conserved viral antigens, whose contribution to influenza immunity are not fully elucidated. To determine the extent to which conserved LAIV antigens contribute to establishing protective immunity against heterologous IAV strains, we generated vesicular stomatitis virus-based single-round vector vaccines expressing individual LAIV antigens, and tested their ability to protect mice from a heterologous challenge with two IAV strains, PR8[H1N1] and rSC35M[H7N7]. We found that immunization with nucleoprotein (NP), ion channel M2, and the stem-region of HA (HA_stem_), but not matrix protein (M1), provide protection from severe disease caused by either IAV strain. This effect correlated with development of non-neutralizing antibodies cross-reactive with both virus strains. Notably, the individual antigens induced specific IgG subclass profiles with different reactivity against PR8 and rSC35M. Sera from vaccinated animals activated Fc-gamma receptor IV-mediated effector functions, suggesting that they can induce cell-mediated immune defense mechanisms, such as antibody-dependent cellular cytotoxicity and antibody-dependent cellular phagocytosis. Combination of the most potent antigens NP and M2 in a mixed vaccination resulted in enhanced protection against IAV challenge, suggesting that the antibody responses against these antigens were synergistic. Our results demonstrate the potency of NP and M2 proteins to serve as conserved antigen targets, resulting in broad protection against severe IAV disease.

## Introduction

Influenza viruses belong to the family of *Orthomyxoviridae*, contain a segmented, single-stranded RNA (ssRNA) genome with negative polarity ^1^, and cause annual epidemics in the human host with 290,000 – 650,000 deaths worldwide ^2^. Of the different influenza types, influenza A virus (IAV) is of most concern for public health, as it induces the most severe pathology in infected individuals, and possesses the highest probability of causing pandemics. In the past, several outbreaks of avian influenza viruses that acquired human-to-human transmissibility through reassortment with human IAV strains illustrate this threat ^3–6^. The viral surface protein hemagglutinin (HA) binds to different cellular sialic acids, predominantly *N*-acetyl-neuraminic acid (Neu5Ac), and thereby initiates the infection of host cells by mediating fusion of the viral and endosomal membrane ^1,7,8^. It is the main target for currently licensed vaccines, which aim to generate virus-neutralizing antibodies (VNAbs) against HA. The VNAb titers generally serve as a correlate of protection ^5,6^. However, since HA is highly variable in its antigenicity, HA targeting vaccines result in suboptimal protection against distinct (mismatched) IAVs. This suboptimal protection promotes the appearance of escape mutants, a mechanism referred to as antigenic drift ^9,10^. IAV vaccines therefore have to be adapted and re-formulated on an annual basis in order to provide sufficient protection against currently circulating IAV strains ^11^. Currently, vaccines commonly in use are mainly inactivated influenza vaccines (IIVs) and live attenuated influenza vaccines (LAIV). IIVs are usually manufactured by inactivating egg- or cell culture-grown virus following disruption of the virus particle using detergents. The HA-standardized vaccine doses are then administered intramuscularly ^11^. In contrast, LAIVs are attenuated, cold-adapted, and temperature-sensitive reassorted virus particles. They harbor internal genes determining the above-mentioned phenotype and surface proteins of a pre-defined IAV strain against which immune responses are directed after vaccination. However, despite the fact that both vaccine preparations contain all viral proteins, they are standardized only regarding their HA content or potency to elicit VNAbs targeting HA. Although more conserved and therefore promising in triggering broad immune responses, internal proteins are understudied in currently applied vaccines, especially in clinical settings.

In the past, many approaches were carried out to broaden or strengthen the immune response after vaccination and circumvent the need for annual re-vaccination and emergence of drifted variants. On the one hand, antigens known to induce a broader antibody response than HA are evaluated for conferring strong, heterologous protection. These include the stem domain of HA (HA_stem_) ^12,13^, the viral neuraminidase (NA) ^14,15^, or the extracellular domain of the ion channel M2 (M2e) ^16–18^. On the other hand, virus-internal proteins, which display a high amino acid conservation between different IAV strains, are a promising target for T cell-mediated immunity. Of most interest are the nucleoprotein (NP) ^19,20^, matrix protein (M1)^21^, and the polymerase subunits, often applied in combination ^22–24^. However, although some approaches proceeded to clinical phases ^25–27^, to date there is no licensed, universal IAV vaccine. Candidates have to overcome major obstacles. As prophylactic agents, they have to be exceedingly safe, compete with currently licensed IAV vaccines, and induce a strong, broad, and long-lasting immunity that can efficiently protect from infection with various IAV strains over several years.

To investigate mechanisms by which broader immune responses can be elicited, we determined the potential of the internal IAV proteins originating from the currently licensed LAIV Fluenz Tetra to induce protection against different subtypes of IAV in mice. We compared these to currently used approaches for universal influenza vaccine like HA_stem_- and M2-based approaches. For immunization, we chose the vesicular stomatitis virus (VSV)-based viral vector platform ^28^. In contrast to, for example, adjuvanted protein vaccine platforms, VSV replicons can be used to express viral proteins in their natural conformation and therefore allow comparison of external surface and internal proteins in their potential to induce cellular, but especially humoral immune responses. VSV-based replicons are attractive vectors for the expression of antigens *in vivo* and are therefore widely used as a vaccine platform against different viruses ^29–32^. One of the main reasons is the low sero-prevalence in the human population, minimizing the chances of vector neutralization by pre-existing immunity. Furthermore, VSV is able to replicate in many tissues by entering the host cells via the widely expressed low-density lipoprotein (LDL) receptor, ensuring optimal infection and subsequent expression of antigens in different cell types ^33^. This leads to high humoral and cellular immune responses. The currently licensed VSV-based vaccine VSV-Ebola virus (VSV-EBOV) demonstrates the feasibility of VSV as a platform to develop effective vaccines ^34^. Deletion of the gene encoding for the VSV glycoprotein (VSV-G) in the VSV-based replicon system (VSVΔG), leads to replication-incompetent vectors, further increasing the safety profile of such approaches ^35,36^.

To compare the protective effect of highly conserved influenza A virus antigens, we generated VSV replicons expressing IAV NP, M1, M2, and HA_stem_ of the live attenuated influenza vaccine (LAIV) Fluenz Tetra and challenged immunized mice with heterologous IAV. Firstly we chose PR8, a mouse-adapted IAV of the subtype H1N1 originating from the A/Puerto Rico/8/34 isolate ^37,38^. Secondly we used rSC35M, a recombinant mouse-adapted H7N7 virus initially emerging during a IAV pandemic in seals (A/Seal/Massachusetts/1/80) that was first passaged in chicken embryo cells and later in mice, resulting in an highly virulent mouse-adapted IAV ^39–41^. We monitored protection from severe illness and lethal outcome and examined underlying T cell-mediated immunity and humoral immune responses including total antibody titers, IgG subclasses, and their ability to induce Fc-gamma receptor (FcγR)-effector functions like antibody-dependent cellular cytotoxicity (ADCC) and antibody-dependent cellular phagocytosis (ADCP). We further immunized mice using a cocktail-vaccination regimen, containing most effective IAV antigens regarding humoral immunity from initial studies and assessed protection and immune responses towards heterologous challenge.

## Results

### Generation and validation of VSV replicon particles expressing IAV proteins

To test the protective potential of conserved proteins against a heterologous IAV challenge, we generated VSV-based single-round replicons expressing individual proteins. As a source we used the currently licensed live-attenuated IAV vaccine Fluenz Tetra (AstraZeneca, season 2017/2018), and cloned open reading frames of NP, M1, and M2 by viral RNA isolation, cDNA synthesis and specific PCR ^42^ (Fig. 1A). Additionally, we generated a HA_stem_-domain construct of the Fluenz Tetra H3 HA (H3_stem_). For this, we substituted the head domain of the HA with a flexible 4x glycine (4xG) linker in order to maintain secondary and tertiary structure of the “headless” H3stem ^43^ (Fig. 1B). The selected antigens displayed a substantially higher amino acid homology to the challenge viruses than the respective full-length HA of the LAIV (Table 1). The respective IAV antigen were inserted into a VSV vector backbone lacking the VSV glycoprotein and expressing GFP (VSV*ΔG; Fig. 1C). In addition to these antigens, a VSV replicon expressing full-length HA protein of the mouse-adapted IAV strain PR8 was used (VSV*ΔG(HA PR8)) ^15^.

**Figure 1:**
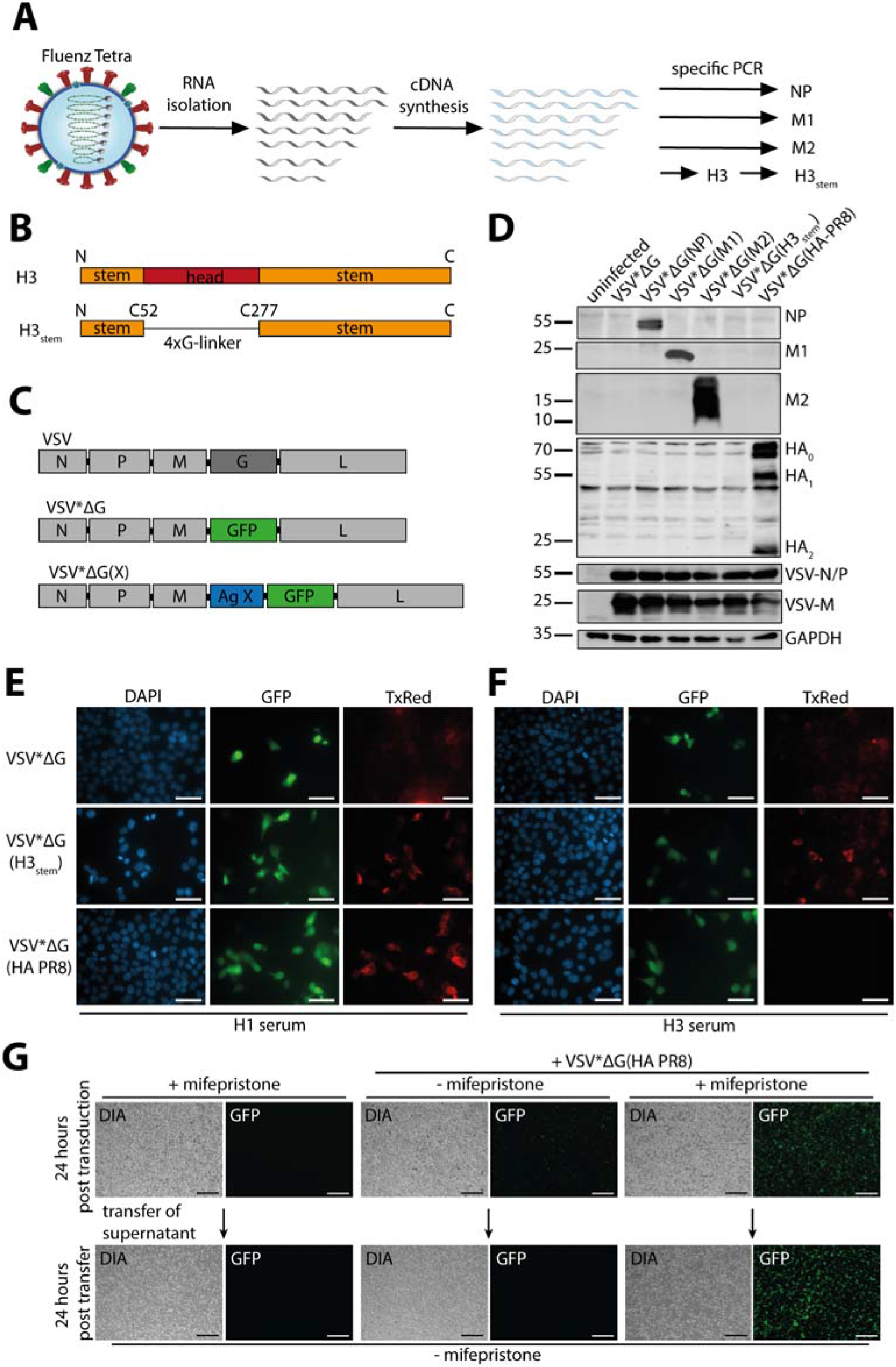
Generation of single-round VSV replicons expressing internal influenza A virus proteins. (A) Graphical illustration of workflow to obtain single influenza A virus (IAV) antigens from the Fluenz Tetra® vaccine virus. RNA was isolated and reverse transcribed. Subsequently, PCR with specific primers was used to amplify genes of interest. (B) Comparison of composition between the full length H3 and the H3_stem_ construct. The full length H3 consists of an N-terminal and a C-terminal part of the stem domain (orange) and the central head domain (red). To receive the H3_stem_ construct, the head domain was replaced with a linker consisting of four glycine residues (4xG-linker, thin line), allowing proper folding of the stem. (C) Genomic organization of VSV (top), VSV*ΔG, in which VSV-G is replaced with GFP (middle), and VSV*ΔG(X) with an additional transcription unit for a foreign antigen X (bottom). (D) Immunoblot analysis to validate the expression of single antigens *in vitro.* BHK-21 cells were transduced with the respective VSV replicons and cell lysates were subjected to SDS-PAGE and immunoblotting for the indicated proteins. (E, F) Immunofluorescence analysis of H3_stem_ and HA expression by VSV replicons. Confluent monolayers of VeroE6 cells were transduced with MOI=1 of the respective VSV replicons. Cells were fixed, and permeabilized 12 hours post infection, and staining was performed using DAPI and either H1 serum (E), or H3 serum (F). Scale bars indicate 50 μm. (G) Verification of single-round replication character VSV*ΔG(HA PR8). BHK-G43 cells were induced to express VSV-G by adding mifepristone and GFP signal was measured 24 hours after transduction with VSV* ΔG(HA PR8) (upper row). To evaluate the release of progeny replicons, supernatant was transferred onto BHK-G43 cells in the absence of mifepristone, and analyzed for GFP signal 24 hours later (bottom row). Scale bars indicate 50 μm.

**Table 1:**
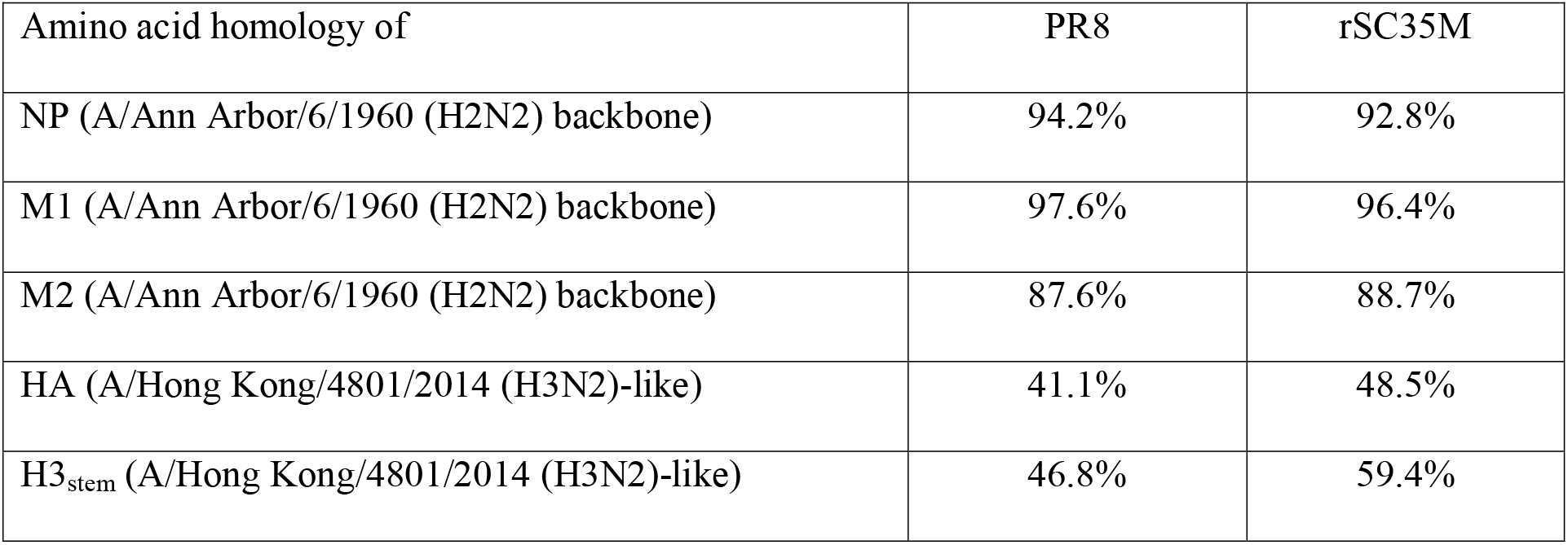
Amino acid homology between antigens expressed in VSV-based replicons and heterologous challenge viruses.

| Amino acid homology of | PR8 | rSC35M |
| --- | --- | --- |
| NP (A/Ann Arbor/6/1960 (H2N2) backbone) | 94.2% | 92.8% |
| M1 (A/Ann Arbor/6/1960 (H2N2) backbone) | 97.6% | 96.4% |
| M2 (A/Ann Arbor/6/1960 (H2N2) backbone) | 87.6% | 88.7% |
| HA (A/Hong Kong/4801/2014 (H3N2)-like) | 41.1% | 48.5% |
| H3<sub>stem</sub> (A/Hong Kong/4801/2014 (H3N2)-like) | 46.8% | 59.4% |

To assess the expression of the different proteins, BHK-21 cells were transduced with the individual VSV replicons, lysed and examined using immunoblot analysis (Fig. 1D). Detection of VSV proteins N, P, and M with a serum obtained from a VSV-immunized rabbit indicated efficient cell transduction with the individual VSV replicons and similar viral protein expression. Specific bands for NP (55 kDa), M1 (25 kDa), and M2 (~15 kDa) demonstrated expression of the individual antigens. In case of the VSV*ΔG(H3stem), no specific band representing H3stem was detected by western blot, whereas expression of full-length HA was confirmed by the characteristic band pattern representing the premature HA_0_ (70 kDa), and the proteolytically cleaved subunits HA_1_ (55 kDa) and HA_2_ (20 kDa) (Fig. 1D).

To provide evidence for expression of H3stem, we performed immunofluorescence microscopy of Vero E6 cells transduced with VSV*ΔG, VSV*ΔG(H3_stem_), or VSV*ΔG(HA PR8). Cells were stained with ferret serum directed against H1 IAV (Fig. 1E), or H3 IAV (Fig. 1F), and a TxRed-conjugated secondary antibody. While GFP^+^ cells were found after transduction with either replicon, a specific TxRed signal was not found in VSV*VG samples. H1 serum, as well as H3 serum, showed reactivity towards the VSV*ΔG(H3stem) transduced cells, as a specific staining occurred on the cell surface of the cells demonstrating efficient expression and trafficking. In contrast to H3 serum, H1 serum was also reactive against VSV*ΔG(HA PR8) transduced cells. In summary, our analyses confirmed expression of NP, M1, and M2, as well as H3stem surface expression with our VSV replicons.

To confirm the single-cycle character of our VSV replicons, we transduced BHK-G43 cells with VSV*ΔG(HA PR8), and either treated the cells with mifepristone or left them untreated (Fig. 1G). Mifepristone induces the expression of VSV-G in BHK-G43 cells ^44^. Increased GFP signal in treated cells indicated efficient replication and propagation, when VSV-G was provided *in trans,* while GFP expression was low in transduced cells that were not treated with mifepristone (Fig. 1G, upper row, compare right and middle panels). Moreover, transfer of supernatants from mifepristone-treated samples to fresh cells led to high GFP-fluorescence, while no GFP signal was detected after transfer of supernatant from untreated cells (Fig. 1G, bottom row), indicating that infectious VSV particles were only released when VSV G was provided *in trans.* Taken together, these results show the successful generation of VSV replicons and expression of the individual IAV antigens upon transduction of BHK-21 cells *in vitro* and confirm the intended single-round character.

### Immunization with VSV replicons expressing IAV NP, M2, and H3_stem_ leads to protection against heterologous IAV challenge in mice

To evaluate the optimal dose for challenge experiments, we titrated the mouse adapted viruses PR8 (H1N1) and rSC35M (H7N7) in C57BL/6J mice leading to a severe, potentially lethal infection (Supp. Fig. 1). PR8 infected mice showed low variation of weight loss in the respective groups (Supp. Fig. 1A). Whereas all animals infected with 1 x 10^2^ TCID_50_ survived the challenge, all higher doses induced a lethal disease (Supp. Fig. 1B). In contrast, rSC35M infected animals exhibited more variable weight loss (Supp. Fig. 1C), and one animal in each group infected with 10^2^ and 10^3^ TCID_50_ survived the infection (Supp. Fig. 1D). We calculated the LD50 from these challenge experiments using the LD50 AAT Bioquest calculator. The LD50 for PR8 was 2 x 10^2^ TCID_50_, and the LD_50_ for rSC35M was 4 x 10^1^ TCID_50_. To trigger severe disease with high mortality, we chose the 3x LD_50_ for all subsequent animal infections (PR8: 6 x 10^2^ TCID_50_; rSC35M: 1.2 x 10^2^ TCID_50_).

To test the protective efficacy of the different IAV antigens, we immunized C57BL/6J mice intramuscularly (i.m.) with 1 x 10^6^ ffu of the respective VSV replicons eight and four weeks before challenge (Fig. 2A). On the day of infection, mice were weighed, anesthetized and infected intranasally (i.n.) with either 6 x 10^2^ TCID_50_ of PR8 (Fig. 2), and we monitored weight loss, disease score, and survival for 14 days (Fig. 2A).

**Figure 2:**
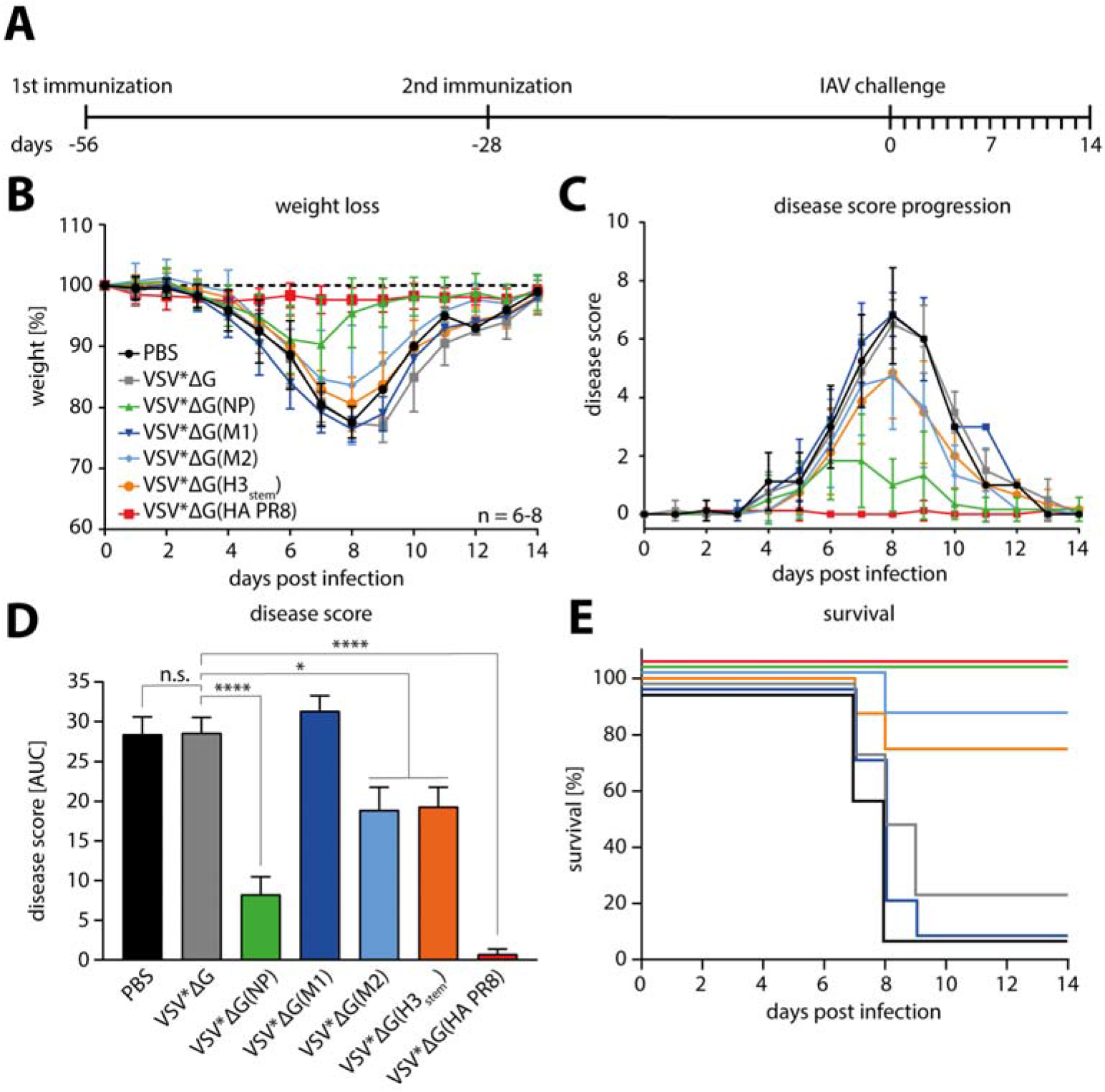
Protection of mice against PR8 challenge after prime-boost immunization with VSV replicons. (A) Schematic illustration of experimental challenge setup. Mice were immunized intramuscularly with 1 x 10^6^ fluorescence forming units (ffu) of the respective replicon 8 weeks (−56 days) and 4 weeks (−28 days) prior to challenge with 6 x 10^2^ TCID_50_ of PR8. After infection, all animals were monitored daily for body weight, behavior, and appearance. (B) Weight loss of PR8 infected animals during the 14 days after challenge relative to their weight on day 0. Dotted line indicates 100%. Each group consisted of n=6 to 8 mice. (C) Disease score of PR8 infected mice representing the sum of numerical values for weight loss, behavior, and appearance. For details, see Supp. Table 2. Data points in (B) and (C) represent means of the respective groups, and error bars indicate standard deviations. (D) Area under curve (AUC) analysis of disease scores. Height of bars represent the mean and error bars are standard error of the mean. For statistical analysis, one-way analysis of variance with Tukey’s multiple comparison post-hoc-test using the VSV*ΔG group as a reference was performed. *p<0.05; ****p<0.0001 (E) Survival analysis of mice infected with PR8.

Mock-immunized mice (PBS), or mice immunized with VSV*ΔG exhibited a severe weight loss of up to 25% after PR8 infection, demonstrating the severe course of infection without pre-existing immunity (Fig. 2B). In sharp contrast, animals immunized with VSV*ΔG(HA PR8) showed no weight loss. This underlines the efficacy of HA-matched IAV vaccines. VSV*ΔG(NP) led to a substantially better outcome in PR8 infected mice in terms of maximal weight loss and recovery time (Supp. Table 1). However, none of the other constructs positively affected body weight loss after PR8 challenge. Since loss of body weight is only one of multiple symptoms of IAV pathology in mice, we determined a more accurate disease score that also accounted for general behavior and appearance of mice in order to assess the clinical course of infection in broader consideration (Fig. 2C, Supp. Table 2) ^45^. While PBS and VSV*ΔG immunized mice showed a severe course of infection, almost no or just mild signs were observed in the VSV*ΔG(HA PR8), or VSV*ΔG(NP) groups, respectively. Interestingly, we also observed lower disease scores in the groups immunized with VSV*ΔG(M2) and VSV*ΔG(H3_stem_), implying protective effects of these vaccines. However, VSV*ΔG(M1) immunization did not show any improvement of the clinical outcome. In order to statistically compare the different disease scores, we calculated the area under the curve (AUC) of disease progression curves (Fig. 2D). Here, we determined a significantly lower overall severity of disease for VSV*ΔG(HA PR8) and VSV*ΔG(NP) (p<0.0001, respectively), and also for VSV*ΔG(M2) and VSV*ΔG(H3stem) (p<0.05, respectively), when compared to VSV*ΔG immunized animals. Assessment of survival reflected these observations by showing low survival rates of mice immunized with PBS, VSV*ΔG, and VSV*ΔG(M1), in contrast to complete survival of VSV*ΔG(NP) and VSV*ΔG(HA PR8) groups, and intermediate survival of VSV*ΔG(M2) and VSV*ΔG(H3_stem_) groups (Fig. 2E).

To test the broad protection of our VSV replicons, we next evaluated their efficacy against a second heterologous virus, rSC35M (Fig. 3). VSV*ΔG(HA PR8) did not show any differences in weight loss (Fig. 3A) or disease score development (Fig. 3B and C) compared to PBS and VSV*ΔG vaccinated control animals. Consequently, this vaccine conferred no positive effect on survival of the mice after rSC35M challenge (Fig. 3D), in contrast to PR8 challenge (Fig. 2E). This underlines the limited efficacy of HA-targeted vaccines against HA-mismatched viruses ^46^. While the VSV*ΔG(H3stem) immunized animals were not protected against weight loss (Fig. 3A), disease scores were slightly decreased (Fig. 3B), although this was tested to be not significant (Fig. 3C). Nevertheless, VSV*ΔG(H3_stem_) vaccination resulted in complete survival of all animals after rSC35M challenge (Fig. 3D). VSV*ΔG(NP) also did not influence development of weight loss (Fig. 3A), but disease scores were significantly lower when compared to VSV*ΔG immunized mice (Fig. 3B and C). However, despite this decreased disease score, not all mice survived the challenge infection of rSC35M (Fig. 3D). Similarly, to VSV*ΔG(H3_stem_), VSV*ΔG(M2) did not decrease weight loss (Fig. 3A), but the overall disease score trended to be lower than in the control groups (Fig. 3B and C), and most animals survived the challenge (Fig. 3D). As observed for PR8 challenge (Fig. 2), VSV*ΔG(M1) immunization did not provide any protection in the rSC35M challenge regarding weight loss (Fig. 3A), disease score (Fig. 3B and C), or survival (Fig. 3D).

**Figure 3:**
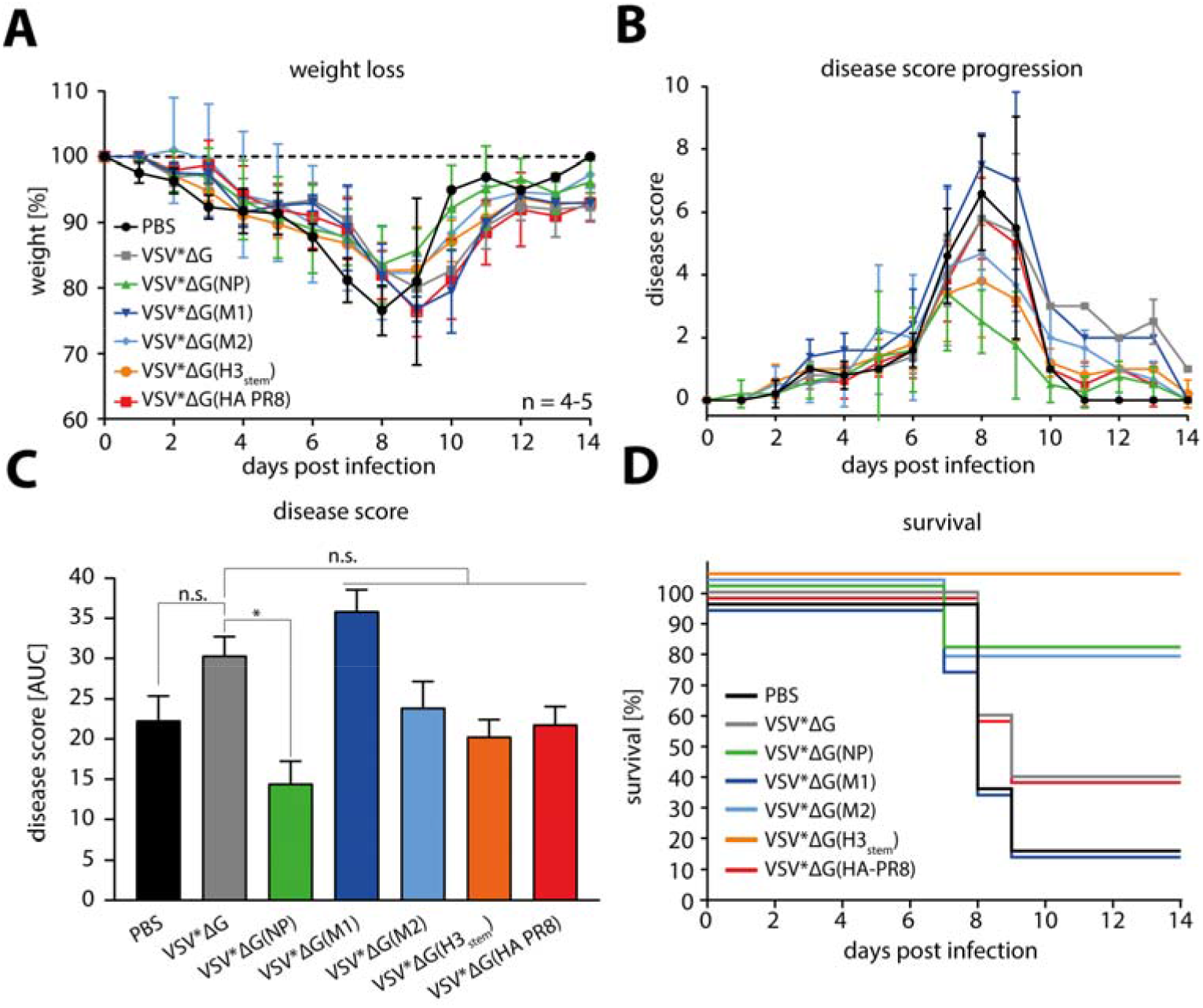
Protection of C57BL/6J mice against rSC35M after prime-boost immunization with VSV replicons. (A) Weight loss of rSC35M infected animals during the 14 days after challenge relative to their weight on day 0. Dotted line indicates 100%. Each group consisted of n=4 to 5 mice. (B) Disease score of rSC35M infected mice representing the sum of numerical values for weight loss, behavior, and appearance. For details, see Supp. Table 2. Data points in (A) and (B) represent means of the respective groups, and error bars indicate standard deviations. (C) Area under curve (AUC) analysis of disease scores. Height of bars represent the mean and error bars are standard error of the mean. For statistical analysis, one-way analysis of variance with Tukey’s multiple comparison post-hoc-test using the VSV*ΔG group as a reference was performed. *p<0.05; ****p<0.0001 (D) Survival analysis of mice infected with rSC35M.

In conclusion, whereas VSV*ΔG(HA PR8) immunization very efficiently protects mice in a homologous challenge model, it was completely inefficient in a heterologous challenge model. While VSV*ΔG(M1) did not improve the clinical outcome of either PR8 or rSC35M challenge, VSV*ΔG(NP), VSV*ΔG(M2) and VSV*ΔG(H3_stem_) immunizations reduced the severity of disease in both severe heterologous challenge models to an extent that improved the survival of the mice.

### Limited T cell responses elicited by VSV replicon immunization

It has been previously shown that individual IAV NP and M1 proteins expressed from a viral vector vaccine can induce IFNγ CD8+ T cell responses in mice ^47^, which could be responsible for the beneficial outcomes in our murine IAV-challenge model. We therefore performed IFNγ and IL-4 ELISpot assays to investigate whether VSV immunization led to generation of antigen-specific T cells that could be activated by either PR8, rSC35M, or Fluenz Tetra viruses (Supp. Fig. 2). In addition, we tested splenocytes of mice infected with low doses of PR8, rSC35M, or Fluenz Tetra for presence of IAV-specific T cells. As virus-specific stimuli, we used a murine dendritic cell line (DC2.4) ^48^ either left uninfected, or infected with PR8, rSC35M, or Fluenz Tetra. Stimulation with PR8 led only to moderate activation of IFNγ-expressing T cells in splenocytes of mice infected with rSC35M, which were the only group of mice exhibiting strong clinical symptoms at the time point of euthanasia. In addition, immunization with VSV*ΔG(NP) led to generation of PR8-reactive T cells in two out of three mice, but this effect was not significant compared to the VSV*ΔG and PBS-immunized control groups (Supp. Fig. 2A). Notably, stimulation with rSC35M-infected DC2.4 led to a strong IFNγ response in all groups independent of the immunization or infection status of the mice, indicating that the active infection with rSC35M caused unspecific IFNγ expression in the assay. Interestingly, stimulation with Fluenz Tetra-infected DC2.4 induced the most robust IFNγ response in splenocytes from mice prior infected with Fluenz Tetra, indicating a limited capability of the vaccine to induce T cell responses against heterologous IAV strains. In contrast to IFNγ, no specific IL-4 T cell responses were observed in any of the groups upon exposure to the various stimuli (Supp. Fig. 2B). From these data we conclude that T cell responses after VSV immunization did not correlate with the protection in the heterologous challenges elicited by the different vectors.

### Total IgG levels correlate with strength of protection against PR8 and rSC35M

We next investigated the humoral immune responses conferred by the different VSV replicons. To do so, we immunized C57BL/6J mice as described above and analyzed IAV-specific antibody titers in the serum four weeks after the first or second dose (Fig. 4A). As expected, none of the groups immunized with either PBS or VSV*ΔG showed antibodies reactive either with PR8 (Fig. 4B) or with rSC35M (Fig. 4C). Analysis of serum from VSV*ΔG(HA PR8) immunized animals revealed high antibody titers against PR8 virus (Fig. 4B, red bars), but not against rSC35M (Fig. 4C, red bars), which is in line with the difference in protective efficacy of this vaccine against the two IAV strains and highlights the narrow humoral immune responses in terms of HA-directed antibodies. In contrast to this, serum from VSV*ΔG(NP) immunized mice had a high reactivity for both PR8 (Fig. 4B, green bars) and rSC35M (Fig. 4C, green bars). Of note, the antibody titers increased in the two-immunizations-regimen. In contrast, only negligible antibody titers were detected in sera from immunizations with VSV*ΔG(M 1). In the VSV*ΔG(M2) group, antibodies were detectable after the second immunization, whereas a single immunization failed to elicit humoral responses. Serum from VSV*ΔG(H3stem) immunized mice showed no reactivity for PR8 (Fig. 4B, orange bars), but high reactivity for rSC35M, at least after prime/boost vaccination (Fig 4C, orange bars). To examine the mechanism of action of these humoral immune responses further, we analyzed the virus-neutralizing activity (Fig. 4D). As expected, no sera showed signs of neutralization of either PR8 or rSC35M, except for the sera from VSV*ΔG(HA PR8) immunized animals, which neutralized PR8, but not rSC35M. These data indicate that immunization with VSV*ΔG(NP), VSV*ΔG(M2), and VSV*ΔG(H3_stem_) induces the generation of IAV-specific cross-reactive antibodies, which may affect disease development after heterologous virus challenge by mechanisms other than virus neutralizing activity.

**Figure 4:**
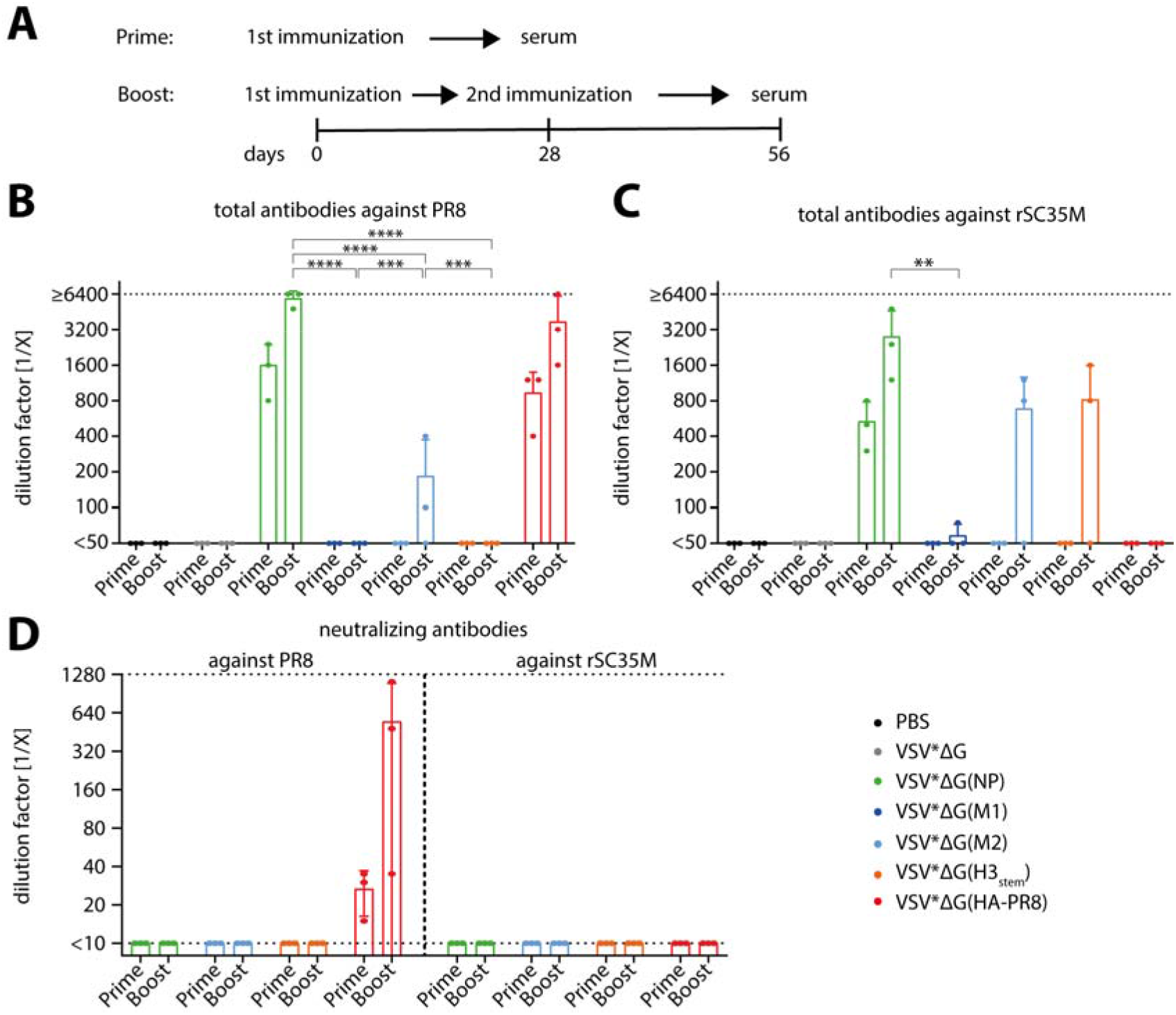
Antibody response of VSV replicon-immunized animals reactive against PR8 and rSC35M. (A) Schematic illustration of the immunization and serum sampling time points. Mice were immunized one or two times and blood was collected either 28 days after prime, or 28 days after boost immunization. (B, C) Total antibody titration after immunization with the respective VSV replicons reactive against PR8 (B) or rSC35M (C). Last wells of infected 96-well plate showing positive staining are expressed as the reciprocal of the serum dilution factor. For statistical analysis, two-way analysis of variance with Dunnett’s multiple comparison post-hoc test was performed on log2-transformed values. **p<0.01; ***p<0.001; ****p<0.0001. Significance levels compared to VSV*ΔG(HA PR8) are not stated. Only significance levels between boost groups of VSV*ΔG(NP), VSV*ΔG(M1), VSV* G(M2), and VSV* G(H3_stem_) are indicated. (D) Neutralizing activity of antibodies against PR8 (left) and rSC35M (right). Last well showing no positive staining is considered positive for virus neutralization and titers are expressed as reciprocals of this dilution. Dotted lines represent the lower and upper limits of detection, respectively. Height of bars indicate mean of groups and error bars are standard deviations.

### VSV replicon immunizations induce different IgG subtype profiles

Mice express several different IgG subclasses (IgG1, IgG2a/c (mouse strain-dependent), IgG2b, and IgG3) ^49^ that mediate different FcγR-effector functions. These effector functions are ADCC, ADCP, and complement activation. IgG1 interacts with the inhibitory FcγR-IIb ^50^, and does not activate complement. The IgG2 subtypes drive ADCC and ADCP by interacting with FcγR-I, -III, and -IV, and can also activate complement via C1q ^51^. IgG3 is a strong complement activator, but it cannot bind FcγRs ^49^. Therefore, the IgG subclass profile can provide insights into the immune response mechanisms elicited by our vaccines. To assess the IgG subclass profiles of mice immunized with the different VSV replicons, we performed a modified immunoperoxidase monolayer assay (IPMA) assay, utilizing secondary antibodies specific for mouse IgG1, IgG2b, IgG2c, or IgG3, respectively (Fig. 5). As before, we quantified the ability of the different IgG subclasses to recognize PR8 infection (Fig. 5A) or rSC35M infection (Fig. 5B). In correlation with total antibodies, VSV*ΔG(NP)-immunized mice showed high PR8-reactive antibody titers of all subclasses, whereas VSV*ΔG(M2) and VSV*ΔG(H3_stem_) mainly generated moderate titers of IgG2b and IgG2c (Fig. 5A). Notably, we observed that some animals failed to generate detectable PR8-reactive antibody levels, making the M2- and H3_stem_-vaccines less consistent than the NP-expressing vaccine. In comparison, the VSV* ΔG(HA PR8) vaccinated group was found to produce high titers of PR8-reactive IgG2b, and IgG2c, whereas IgG1 expression was inconsistent among the animals in this group, and IgG3 was not detectable (Fig. 5A).

**Figure 5:**
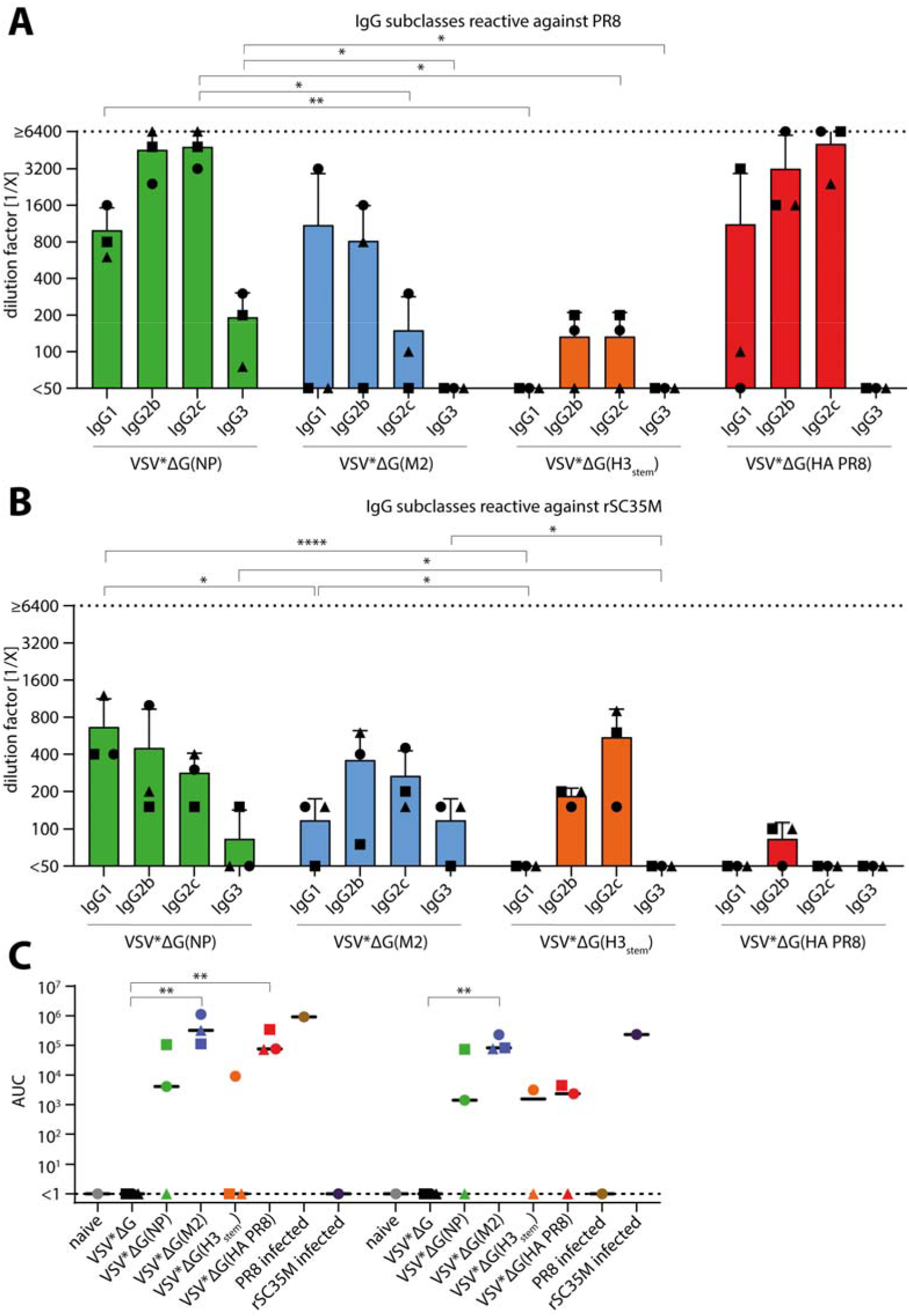
IgG subclasses and FcγR-IV activation. (A, B) IgG subtype analysis of immune sera 28 days after boost vaccination against PR8 (A) and rSC35M (B). Only serum with detectable total antibodies were subjected to analysis. Bars represent the mean, and error bars are standard deviations. Dotted lines represent the lower and upper limits of detection, respectively. Last wells showing positive staining are expressed as the reciprocal of the serum dilution factor. For statistical analysis, one-way analysis of variance with Tukey’s multiple comparison post-test was used, and all groups were compared. Levels of significance are shown accordingly; *p<0.05; **p<0.01; ***p<0.001. (C) PR8- or rSC35M-infected MDCK cells were incubated with 2-fold dilutions of serum, and FcγR-IV expressing reporter cells were added. Luminescence was measured and area under curve (AUC) was calculated. Serum of a naïve mouse plus three times its standard deviation was used as a baseline. Symbols indicate sera from individual animals, and black bars illustrate mean values in each group. For statistical analysis, two-way analysis of variance with Sidák’s multiple comparison test on log2-transformed titers was conducted. Dotted line indicates lower limit of detection.

With few exceptions, IgG subtype titers reactive against rSC35M (Fig. 5B) followed a similar trend as seen for the PR8-reactive antibodies, but the elicited rSC35M-reactive antibody titers were generally lower than the PR8-reactive antibodies. VSV*ΔG(NP) vaccination induced similar levels of IgG1 and IgG2 against rSC35M, and low levels of IgG3. In contrast, VSV*ΔG(M2) and VSV*ΔG(H3_stem_) vaccination led to higher IgG2b and IgG2c titers than IgG1 or IgG3. Notably, VSV*ΔG(HA PR8)-immunized mice had no or very low rSC35M-reactive IgG of all subclasses, even though these sera contained high titers of PR8-reactive IgG. In conclusion, IgG subtype analysis revealed that VSV*ΔG(NP)-vaccination induced the broadest profile of IgG subclasses reactive against both heterologous IAV strains. In addition, the M2- and H3stem-expressing vectors induced mainly IgG2 subclasses, which were also reactive with both PR8 and rSC35M, albeit these antibodies were not consistently generated in all vaccinated animals.

### Antibodies directed against NP, M2, and H3stem can activate FcγR-IV

To evaluate mechanisms of protection with this strategy, we first investigated potential FcγR-mediated effector functions (Fig. 5C). Since IgG2b and IgG2c titers were consistently higher than IgG1 and IgG3 titers for our vaccine constructs, we utilized a murine FcγR-IV reporter assay, since this receptor is almost exclusively activated by IgG2 subtypes ^49,52^. For this, IAV-infected MDCK cells were co-incubated with the respective sera and reporter cells expressing FcγR-IV, and a luciferase reporter driven by a nuclear factor of activated T cell (NFAT)-response element. Serum of VSV*ΔG(M2)-immunized animals efficiently activated FcγR-IV in the presence of either PR8 or rSC35M infected cells (Fig. 5C). We also observed strong FcγR-IV activation by sera from VSV*ΔG(NP) and VSV*ΔG(H3stem) groups, although not all sera in these groups activated FcγR-IV consistently. Antibodies after VSV*ΔG(HA PR8) immunization activated FcγRIV after incubation with PR8 infected cells more efficiently than after incubation with rSC35M infected cells. Control sera from mice infected with either PR8 or rSC35M only activated FcγR-IV when incubated with cells infected with the homologous virus, but not with the heterologous virus. Taken together, our results provide evidence that non-neutralizing antibodies directed against NP and M2 can activate FcγR effector functions, and thereby contribute to protection against heterologous IAV challenge.

### Synergistic effects of NP- and M2-expressing VSV replicons on protection against PR8 challenge

Based on the observation that single VSV*ΔG(NP) immunization led to strongest protection while VSV*ΔG(M2) vaccination resulted in highest FcγR-IV activation, we hypothesized that the two antigens provide immunization by different, potentially synergistic mechanisms. In order to examine this complementary potential, we vaccinated mice with both VSV replicons at once. For this, we used a cocktail of VSV*ΔG(NP) + VSV*ΔG(M2) (double-cocktail) using 5 x 10^5^ ffu of each replicon, and compared this to 5 x 10^5^ ffu of only NP or M2, respectively, complemented by VSV*ΔG to obtain a total dose of 1 x 10^6^ ffu of replicons for all approaches (Fig. 6). We used the same prime-boost and challenge experimental setup as described for prior experiments (Fig. 2A). As expected, empty vector-immunized animals (VSV*ΔG) showed severe pathology after challenge with PR8, with massive weight loss of up to 25% on average (Fig. 6A), high disease score (Fig. 6B and C) and a majority of mice succumbing to disease (Fig. 6D). Interestingly, vaccination with the VSV double-cocktail (0.5x VSV*ΔG(NP) + 0.5x VSV*ΔG(M2)) resulted in complete protection against PR8-induced pathology in terms of weight loss (Fig. 6A), significantly (p<0.001) decreased disease score (Fig 6B and C) and led to 100% survival (Fig. 6D). Notably, the double-cocktail was more efficient that either low dose of the individual VSV-replicons, which suggests a complementary protection via NP- and M2-directed immune responses. However, even half the dose of initially injected VSV*ΔG(NP) led to protection, demonstrated by significant (p<0.001) reduction of overall disease score (Fig. 6C) and 100% survival (Fig. 6D). In parallel, we further investigated the effect of the three most promising antigen candidates for protection NP, M2, and H3_stem_ in a triple-cocktail approach (Supp. Fig. 3). For this, we mixed 1/3 doses of VSV*ΔG(NP), VSV*ΔG(M2), and VSV*ΔG(H3stem), and compared protection to 0.3x VSV*ΔG(NP) and 0.3x VSV*ΔG(M2) using the established immunization-challenge protocol (Fig. 2A). Surprisingly, when three antigens were combined in one triple-cocktail vaccination, protection against heterologous PR8 infection was inferior to double-cocktail as demonstrated by striking weight loss of up to 15% (Supp. Fig. 3A), intermediate disease scores (p<0.05) (Supp. Fig. 3B and C), and reduced survival rates (Supp. Fig. 3D).

**Figure 6:**
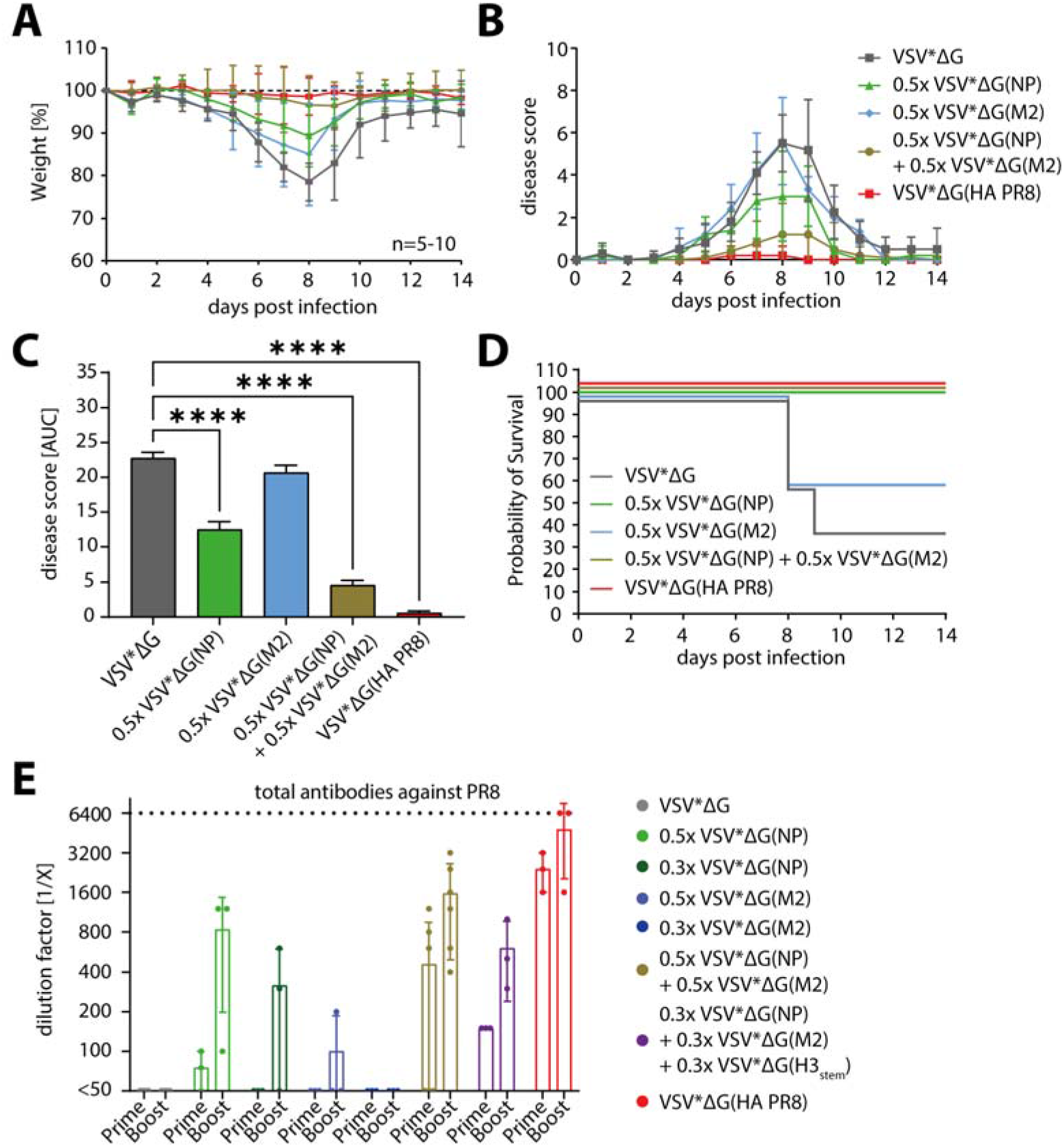
Protection of C57BL/6J mice after prime-boost immunization with VSV replicon double-cocktail immunizations against PR8. (A) Weight loss of PR8 infected animals during the 14 days after challenge relative to their weight on day 0. Dotted line indicates 100%. Each group consisted of n=5 to 10 mice. (B) Disease score of PR8 infected mice representing the sum of numerical values for weight loss, behavior, and appearance. For details, see Supp. Table 2. Data points in (A) and (B) represent means of the respective groups, and error bars indicate standard deviations. (C) Area under curve (AUC) analysis of disease scores. Height of bars represent the mean and error bars are standard error of the mean. For statistical analysis, one-way analysis of variance (ANOVA) with Tukey’s multiple comparison post-hoc-test using the VSV*ΔG as a reference was performed; ****p<0.0001. (D) Survival analysis of mice infected with PR8. Surviving mice were euthanized at the end of the study. (E) Total antibody titration against PR8. Last wells showing positive staining are expressed as the reciprocal of the serum dilution factor. Dotted line represent the upper limit of detection. Height of bars indicate mean of groups and error bars are standard deviations.

To investigate the underlying humoral immune responses after cocktail vaccination, we analyzed the serum of immunized animals and determined titers of total IgG (Fig. 6E). Total antibody titers correlated with protection during challenge experiments, as PR8-specific titers were higher, although not significant, when animals were immunized with the double-cocktail as compared to the triple-cocktail (Fig. 6E). In line with our expectations based on prior experiments (Fig. 4B), we observed highest titers among individual replicon vaccinations for 0.5x VSV* ΔG(NP), and only weak titers for 0.5x VSV*ΔG(M2). These were even decreased after further replicon dose dilution, as demonstrated for 0.3x VSV*ΔG(NP) and 0.3x VSV*ΔG(M2), respectively. To examine the induced IgG subclass profiles further, we performed an IPMA specific for either IgG subtype for cocktail immunized mice. Interestingly, no differences for either virus or subtype was detected, as intermediate IgG1, high IgG2b and IgG2c and low IgG3 levels were observed throughout (Supp. Fig. 3E-H).

Finally, we investigated the effect of the double- as well as triple-vaccination approaches also on challenge infection with rSC35M (Supp. Fig. 4). Here, both cocktail immunizations failed to protect rSC35M-challenged mice from severe weight loss (Supp. Fig. 4A), and animals of all groups developed high disease scores (Supp. Fig. 4B). In terms of survival, double- and triple-cocktail immunizations protected 40% of animals from death (Supp. Fig. 4C). While this was lower than the 100% and 80% survival achieved in the PR8-challenge (Fig. 6D and Supp. Fig. 3D), it was still slightly better than the vaccination with the mismatched HA PR8-vector, in which group only 20% of mice survived the rSC35M-challenge (Supp. Fig. 4C). Interestingly, despite the weak protection of the double- and triple-cocktail immunizations in the rSC35M challenge, high levels of total antibodies reactive towards rSC35M (Supp. Fig. 4D), as well as IgG subclass titers were observed (Supp. Fig. 4E-H), suggesting that high levels of non-neutralizing antibodies alone do not always protect from developing severe disease.

In conclusion, our results underline that internal proteins of IAV can be used as an immunogenic target, as they can induce strong humoral immune responses that are able to confer protection in the absence of neutralization. Activation of FcγR-ΓV effector functions strongly suggest ADCP and ADCC as protecting mechanisms, which lead to beneficial outcome after heterologous IAV challenge in terms of disease severity and overall survival. This underlines the protective potential of non-neutralizing antibodies and can serve as a basis for the rational design of future heterologous IAV vaccines.

## Discussion

Currently licensed IAV vaccines provide sub-optimal protection against infection and drive antigenic drift of seasonal influenza virus ^9^. Furthermore, emergence of different avian IAVs that crossed the species-barrier in the past fuel the fear of potential pandemics of new reassortants ^53–55^. Since seasonal influenza vaccines are adapted to the currently circulating strains, it is highly probable that these vaccines would fail to protect efficiently from emerging IAV. It is therefore urgently needed to refine these vaccines in terms of strength, breath, and duration of the elicited, protective immunity. In the past, different approaches were used to achieve this enhancement, manly by targeting conserved antibody domains like the stem region of HA (HA_stem_) or the ectodomain of the M2 ion channel (M2e) ^12,16^, or by inducing cytotoxic T cell responses towards internal proteins like NP or M1 ^19,21^. Using VSV replicons to express the proteins NP, M1, M2, and H3_stem_ originating from the LAIV Fluenz Tetra, we compared their potential to induce protective immune responses and analyzed underlying immunity towards these antigens. We show that NP, M2, and H3_stem_-expressing vectors, in contrast to M1, protect mice from severe PR8 infection, and to some degree from rSC35M infection, and that this protection correlates with total antibody levels and IgG2 subclass titers. In line with this, we demonstrated FcγR-IV activation after NP and especially after H3_stem_ immunization, suggesting ADCC and ADCP as mechanisms of protection. Remarkably, by combining VSV*ΔG(NP) and VSV*ΔG(M2) in a double-cocktail immunization approach, synergistic effects were observed. Our results highlight the potential of especially the internal protein NP to not only serve as an T cell target, as plentifully described in the past ^23,24,56–59^, but also as a target for humoral immunity. These results can help better understand the underlying mechanisms of non-neutralizing antibodies against highly conserved IAV antigens to provide heterologous protection and nourish the rational design of universal influenza vaccines in the future.

### Protection against heterologous IAV challenge after VSV replicon immunization correlates with humoral immune responses towards internal proteins

Conserved epitopes of IAV have been a major target for the development of universal influenza vaccines in the past. Although enormous effort has been made to elicit protective cellular immunity towards these antigens by a variety of immunization approaches, to date there is no licensed IAV vaccine targeting those ^60–62^. In contrast, humoral immune responses against internal, conserved IAV proteins were neglected, as they are thought to be less accessible for antibodies, and conferred protection is expected to be inferior to surface antigens, mostly due to the lack of neutralization activity. However, it was shown that the internal protein NP can be presented on the surface of infected and dying cells, and that monoclonal antibodies against NP can protect against IAV pathogenicity and high viral lung titers in the mouse model, when expressed *in vivo* or even after serum transfer ^63–66^. We show that in our mouse model, VSV replicons are able to induce antibodies against internal (NP) and external (M2, H3_stem_) proteins, which is in line with previous studies ^47^. However, VSV*ΔG(M1) failed to do so. Protection against the potentially lethal doses of PR8 or rSC35M correlated with the levels of total antibodies against the various antigens. Importantly, while VSV*ΔG(NP) and VSV*ΔG(HA PR8) induced comparably high humoral responses protecting against PR8 virus, only NP-induced antibodies interacted with the heterologous rSC35M. This is in line with studies showing the potential of VSV-vectored vaccines expressing IAV NP as an antigen to elicit partially protective immune responses ^67^. This underlines the major drawback of full HA-based vaccines. Remarkably, our construct representing the widely used HA_stem_ domain targeting approaches, led to humoral reactivity towards rSC35M, but not PR8. HA can be categorized into two different groups, group 1 HA and group 2 HA, based on their phylogenetic similarities ^68^. While H1 (PR8) belongs to group 1 HAs, H3 (Hong Kong-like) and H7 (rSC35M) are classified in group 2 HA. This in in line with our amino acid homology analysis, showing a higher similarity between H3 and H7 (59.4%) than between H3 and H1 (46.8%) and stresses the challenge of HA_stem_-based approaches, namely the broad protection against both groups of HA (Table 1). This observation, in combination with the fact that also VSV*ΔG(NP) immunization protected better against PR8 (94.2% homology) than rSC35M (92.8% homology), suggests that immunity against conserved antigens can provide heterologous protection, but its degree may depend on the level of homology. For the rational design of future heterologous IAV vaccines, we therefore hypothesize that it is necessary to include multiple IAV antigens.

The proposed mechanisms of action for non-neutralizing antibodies are based on FcγR-effector functions, and include ADCC, ADCP, and complement activation ^69^. Mice express four subclasses of IgG that differ in their FcγR-effector functions. For example, while IgG1 cannot activate complement and inhibits IgG2 binding to FcγR ^70^, IgG3 activates complement very efficiently, but does not interact with FcγRs ^49^. IgG2c and IgG2b activate both pathways and may represent a correlate of protection in terms of efficient ADCC or ADCP activation. However, the complex interplay between the different IgG subclasses is not fully understood and more insights are needed to resolve the different, overlapping, or subsidiary functions of murine IgG reliably. Future experiments investigating complement activation in the lung may complement our current understanding of protective, non-neutralizing antibody responses in the presented setting.

It is well-known that the modality used to immunize mice is crucial for the shape of the IgG subclass expression ^71–73^. Our IgG subclass analysis revealed significant differences concerning the humoral immune response induced against the single IAV antigens and most remarkably is the strikingly high IgG2c and IgG3 titer of VSV*ΔG(NP) immunized animals when compared to the other groups, a phenomenon also observed in IAV infected mice ^71,74^. As VSV*ΔG(NP) groups showed a lower level of weight loss and a faster recovery during challenge experiments, IgG2c and IgG3 titers correlated with protection. Additionally, low titers of IgG1 were detected. As a recent publication found that not-neutralizing antibodies of the IgG1 subclass can suppress IgG2-driven effector functions, this strengthens the expectation of a highly protective IgG subclass profile ^75^. To investigate this, we decided to measure murine FcγR-IV activation as this receptor is known to not only induce ADCC and ADCP, but also to have a high affinity and selectivity for IgG2b and IgG2c subclasses ^52^. We therefore hypothesized that this approach best translates the observed IgG2b/c subclass profiles into the Fc-mediated effector functions. Asthagiri Arunkumar et al. used reporter cells expressing the human FcγR-IIa, which is known to cross react with all murine IgG subclasses except for IgG3, and found ADCC after immunization with a viral MV A vector expressing NP and M1 of IAV ^47,52^. Nevertheless, the approach of measuring murine IgG-induced Fc-mediated effects by human receptors is questioned by a recent publication finding relatively weak interaction between those ^76^. Finally, both experiments utilizing reporter cells expressing either human FcγR-IIa or murine FcγR-IV showed that non-neutralizing antibodies directed against internal proteins of IAV can in fact induce Fc-mediated effector functions against heterologous viruses. Strikingly, when we tested the serum of a control animal infected with either PR8 or rSC35M, we only detected strain specific activation of murine FcγR-IV, suggesting that the immunodominance of HA during IAV infection outweigh other epitopes. Remarkably, against both viruses, M2-directed antibodies of all animals had a high capability to activate FcγR-IV. This in in line with current understanding of M2-directed antibodies as they are not neutralizing but known to be protective in the murine model ^77^. Lack of protection via M2e-specific antibodies in FcγR-/- mice highlights this necessity for Fc-mediated effector functions and that a dominant mechanism is ADCC by alveolar macrophages expressing FcγR-IV among others ^78,79^. On average, we found higher FcγR-IV activation in NP immunized animals when compared to the H3stem group against both viruses. However, besides neutralizing antibodies directed against the stem region of HA, which we did not detect for either virus, ADCC and complement activation through antibodies are considered to be major drivers of protection ^80,81^. FcγR-IV activation in NP-immunized animals against PR8 and rSC35M was comparable, which contradicts the concept of IgG2-based immunity supported by the different IgG subclass profiles. However, our findings underline the potential of NP and M2 to induce non-neutralizing, FcγR-IV-activating humoral responses against heterologous viruses after challenge. We chose a restrictive approach by utilizing FcγR-IV-expressing reporter cells. Future experiments using, for example, FcγR-IIB and FcγR-III could evaluate if the Fc-mediated effector functions are indeed even stronger and to what extent IgG1 is important in this regard. This is especially important as many immune cells, for example NK cells, express FcγR-IIB and FcγR-III, which are strong drivers of ADCC ^52^

The faster recovery and limited weight loss of VSV*ΔG(NP) vaccinated mice is characteristic for non-sterile immunity, often attributed to T cell immunity. As NP can be a strong driver of T cell immunity, we first hypothesized that the protection in our immunization-challenge model is based on cellular immunity ^22,82,83,83–85^. However, we detected only limited increases in IFNγ, and no increase in IL-4 secreting T cells after immunization with our various VSV replicons. Possible explanations include the choice of the vector to deliver the IAV antigen, as it was shown in a comparable study that immunization with VSV*ΔG(X) replicons expressing IAV neuraminidase did not lead to measurable cellular immunity ^15^. However, another study reported strong CD8+ T cells after immunization in mice using a VSV vector expressing IAV NP ^67^. Remarkably, in this study homologous PR8 NP immunization and ELISpot stimulation with the immunodominant epitope NP_366-374_ were used. However, this epitope differs between the NP of LAIV, which was used for immunization here, and the NP of the challenge viruses PR8 or rSC35M, which may explain the limited efficacy of our vector to induce T cell responses. Whether T cell or humoral responses were the major drivers of protection, or both partially contributed to the efficacy of our replicon immunizations, has to be clarified in future studies using adoptive T cell and serum transfers.

### Combination of most promising antigens leads to variable outcomes against PR8 and rSC35M challenge

Although previous studies have demonstrated that single external, as well as internal, antigens can induce at least partial protection against IAV challenge in mice ^62,86–88^, it has become increasingly clear that future vaccines most probably have to combine more than one IAV antigen. We therefore chose our most promising VSV replicons from initial challenge experiments and tested conferred protection via double- (VSV*ΔG(NP) and VSV*ΔG(M2)) and triple- (VSV*ΔG(NP) and VSV*ΔG(M2) + VSV*ΔG(H3_stem_)) cocktail immunizations. Interestingly, the double cocktail induced highly protective immune responses against PR8 challenge. In this group, mice experienced no weight loss or development of any symptoms at all, indicating synergism between the immune responses elicited against NP and M2 antigens. While FcγR-IV activation was strongest for VSV*ΔG(M2)-immunized animals, total and IgG subclass titers were shown to be moderate. In contrast, VSV*ΔG(NP) total as well as IgG2/3 titers were high, but FcγR-IV activation was intermediate. In this model, M2 directed antibodies may induce ADCC against infected and living cells more efficiently than NP directed antibodies, while NP directed antibodies may preferentially recognize apoptotic cells, and thereby enhance ADCP and subsequent antigen presentation, which in turn may leverage adaptive immunity. Future studies should address and differentiate the exact Fc-effector mechanisms induced by anti-M2 or anti-NP antibodies. However, our results clearly underline the complementary principle of multiple antigens used in many approaches ^89^. In contrast to the double-cocktail, the triple-cocktail generated inferior protective effects. It has to be stressed that the total dose of replicons during all immunizations was 1 x 10^6^ ffu/mouse. Animals vaccinated with the double- or the triple-cocktail therefore only received a reduced dose of each VSV vector, including the most protective VSV*ΔG(NP). Studies in mice immunized with varying doses of IAV vaccines demonstrated the significant differences in produced total and subclass antibody titers ^90,91^. As it is known for influenza vaccines to induce a more robust immune responses when administered in higher doses, a possible explanation may be that by diluting VSV*ΔG(NP) we undercut the threshold of antigen needed to elicit strong responses for the triple-cocktail ^92,93^.

Surprisingly, in rSC35M infected animals, neither the double-cocktail nor the triple-cocktail immunization had a beneficial effect, even though we found rSC35M-detecting total and subclass-specific IgG titers in these sera to be in the same range as their PR8-detecting counterparts. While a mechanistic explanation will require further analyses, it is conceivable that in the pool of antibodies generated in response to the VSV replicon vaccinations, only certain antibody clones efficiently mediate mechanisms such as ADCC and ADCP. These antibodies may have varying affinity for the antigens of the two challenge viruses used here, or lower rSC35M-specific than PR8-specific titers, which could be masked by high levels of antigen-recognizing, but effector-inefficient antibodies in case of rSC35M.

Overall, our study demonstrates the high potential of internal IAV proteins, especially NP, to induce humoral immune responses that can contribute to protection after a potentially lethal challenge with a heterologous IAV. This protection is further optimized by adding M2, indicating synergistic effects of NP- and M2-directed immune responses. However, variable degrees of efficacy of our approach against heterologous IAV strains highlight the complex nature of immune responses induced by the same vaccine approach against different viruses. It is therefore advisable to further investigate and implement internal proteins as a target for humoral immunity in the rational design of future IAV vaccines.

## Materials and Methods

### Cell culture

Baby hamster kidney cells (BHK-21) cells were maintained in Eagle’s minimal essential medium (MEM; Sigma Aldrich) supplemented with 10% fetal bovine serum (FBS; Life Technologies) and 200 mM L-glutamine (Sigma-Aldrich) at 37°C and 5% CO_2_. For BHK-G43 ^94^ cells, which express the VSV-G protein upon mifepristone induction, 0.5 mg/mL Zeocin (Invitrogen) and 125 μg/mL Hygromycin B (Sigma-Aldrich) were added every third cell passage. Madin-Darby Canine Kidney (MDCK) and VeroE6 cells were maintained in Dulbeccos’s modified Eagle medium (DMEM; Sigma-Aldrich) supplemented with 5% FBS, 200 mM L-glutamine and 100 U/mL penicillin/streptomycin (Life Technologies).

### Influenza A virus production

Mouse-adapted influenza virus A/Puerto Rico/8/1934(H1N1) (PR8) and recombinant mouse-adapted A/Seal/Massachusetts/1-SC35M/1980(H7N7) (rSC35M) ^41^ (kind gift from Prof. Dr. Friebertshäuser; Philipps-University Marburg) were grown in eleven days old embryonated chicken eggs. For this, 1 x 10^3^ TCID_50_ of virus was inoculated into the allantoic fluid and eggs were incubated for 1-2 days at 37°C and 60 % humidity in an egg incubator. Allantoic fluid was harvested by a pipette into a falcon tube and centrifuged for 5 min at 500 g and 4°C. Hemagglutination titers of the supernatant were determined as follows: Supernatants were two-fold diluted in a U-shaped 96-well plate and 50 μL of 1% turkey-erythrocyte-suspension was added. After 30 min at 4°C, hemagglutinin titers were determined according to standard protocols ^95^ and supernatants with comparable titers where pooled. Stocks were aliquoted and frozen at −80°C. For cell culture-grown virus stocks, MDCK cells were infected with either PR8 or rSC35M in serum-free DMEM supplemented with tosylsulfonyl phenylalanyl chloromethyl ketone-treated Trypsin (TPCK-trypsin; Sigma-Aldrich) with a concentration of 0.75 μg/mL. Cells were observed daily and supernatant was harvested and centrifuged for 15 min at 3000 g and 4°C as soon as cytopathic effect was widespread. Cleared supernatants were stored at −80°C.

### Influenza A virus titration

For titration of virus, 2 x 10^6^ MDCK cells were seeded in flat-bottom 96-well plates and incubated for one day at 37°C and 5 % CO_2_ to reach 90 % confluence. 10-fold dilutions of virus were added in quadruplicates and incubated for additional two days at 37°C and 5 % CO2. Plates were washed with 1:3 PBS: double-distilled H2O (ddH2O), and air-dried for 20 minutes prior to heat fixation at 65°C for 8 hours. Plates were then incubated with 50 μL of ferret anti-PR8 serum (Paul-Ehrlich-Institute) diluted 1:500 in PBS, or mouse anti-Influenza NP (Cat#: MA5-29926; Invitrogen) diluted 1:2,000 in PBS, for 2 hours at room temperature. Plates were washed once for 20 min with 100 μL PBS and subsequently incubated for 1 hour at room temperature with either goat anti-ferret IgG-h+l horseradish peroxidase (HRP)-conjugated antibody (Cat#: A140-108P; Bethyl Laboratories), or goat anti-mouse HRP-conjugated antibody (Cat#: 115-035-003; Jackson Immunoresearch), diluted 1:750 in PBS. After incubation, plates were washed once for 20 min with PBS, and 3-amino-9-ethyl-carbazol (AEC) substrate (Sigma Aldrich) diluted in N,N-Dimethylformamide (DMF) (Cat#: D4451; Sigma Aldrich) was added. After visible staining occurred, substrate was discarded and wells were washed with 100 μL of ddH_2_O to stop the reaction. Positive wells were visualized by red staining of infected cells and were expressed as the 50 % tissue culture infectious dose per milliliter (TCID_50_/mL).

### Generation of VSV replicon cDNA

For all experiments, we used VSV vectors expressing green fluorescent protein (GFP; indicated by *) and lacking VSV-G (ΔG). VSV*ΔG and VSV* ΔG(HA PR8) were previously generated ^15^. To generate the remaining VSV replicons, we isolated viral RNA directly from the Fluenz Tetra vaccine (AstraZeneca; season 2017/2018) virus using the Qiagen Viral RNA Isolation Kit (Qiagen) following the manufacturer’s instructions. The Uni12 primer ^42^ was used for reverse transcription in order to obtain viral cDNA, and segment-specific primer pairs including restriction sites for subcloning were subsequently used for amplification of the respective open reading frames encoding the NP, M1, and M2 proteins (Supp. Table 3). To generate the H3stem construct, overlapping primers of the two distinct stem regions were utilized to receive a headless HA gene by subsequent joining PCR. The subcloned ORFs were then inserted into an expression plasmid encoding the VSV*ΔG full-length genome using restriction sites *MluI* and *BstEII.* Sequence identities were confirmed by Sanger sequencing (Eurofins Genomics).

### VSV replicon rescue and titration

To generate VSV replicons, BHK-G43 cells treated with 1 nM mifepristone to induce expression of VSV-G and subsequently transduced with a modified vaccinia virus Ankara encoding for the T7 phage RNA polymerase (MVA-T7) ^96^. Cells were then co-transfected with VSV helper plasmids (VSV-N, -P, -L) and the respective genomic plasmid, in which the VSV genome is controlled by the T7 promotor, as previously described ^94^. 24 hours after transfection, BHK-43 cells were trypsinized, seeded together with an equal number of fresh BHG-43 cells, and incubated with mifepristone for another 24 hours ^97^. Replicon-containing supernatant was then centrifuged for 15 minutes at 1,000 g and 4°C, aliquoted and stored at −80°C. To produce VSV replicon stocks suited for animal immunization, BHK-G43 cells seeded to reach 90 % confluence and VSV-G expression was induced by adding 1 nM mifepristone for 6 hours. VSV replicons were added. Supernatant was harvested as soon as GFP expression was widespread (approx. 48 hpi), and centrifuged for 15 min at 1,000 g and 4°C. VSV replicon-containing supernatants were concentrated by centrifugation through a 20 % (w/v) sucrose cushion, for 1 hour at 100,000 g and 4°C in a SW-32Ti rotor. Afterwards, replicon pellets were resuspended in sterile PBS, and stored at −80°C. For titration of VSV replicons, 2 x 10^4^ BHK-21 cells were seeded in 96-well plates to reach 90 % confluence the next day. After 24 hours, VSV replicons were applied in 40 μL of two-fold dilutions in duplicates and incubated for 90 minutes at 37°C and 5% CO2. Subsequently, 60 μL of MEM medium were added, and cells were incubated overnight. To determine replicon titers, GFP-positive cells were counted in an appropriate dilution and titers were calculated and expressed as fluorescence-forming units per mL (ffu/mL).

### Immunoblot analysis

For immunoblot analysis, confluent BHK-21 monolayers were transduced with the respective VSV replicon in MEM medium at 37°C. After 24 hours, cells were lysed, and lysates were subjected to immunoblot analysis as previously described ^98^. Primary antibodies were diluted in TBS supplemented with 0.1% -Tween-20 and 2.5% BSA at the indicated dilutions. Primary antibodies used were goat anti-Influenza A virus (1:200; Paul-Ehrlich-Institute), mouse anti-Influenza A Virus M1 (1:1,000; clone: GA2B; Cat#: MA1-80736; Invitrogen), mouse anti-Influenza A virus M2 (1:1,000; clone: 14C2; Cat#: MA1-082; Invitrogen), mouse anti-GAPDH (1:1,000; Cat#: AKR-001, Cell Biolabs Inc.), rabbit-anti Influenza A H1N1 HA (1:1,000; Cat#: PA5-81670; Invitrogen), rabbit anti-VSV (1:5,000; Prof. Dr. Conzelmann, Gene Center Munich) ^32^. Secondary antibodies, prepared in TBS-Tween at the indicated dilutions, were: bovine α-goat IgG-HRP (1:20,000; Cat#: sc-2384; Santa Cruz Biotechnology), goat α-rabbit IgG Fc conjugated to horseradish peroxidase (HRP) (1:20,000; Abcam), goat α-mouse IgG(H+L)-HRP (1:20,000; Jackson Immunoresearch). Chemiluminescence detection was performed using the Amersham ECL Prime Western Blotting Detection System (Cytiva Amersham™) and a ChemiDoc Imager (Bio-Rad) with ImageLab software (version 6.0.1).

### Immunofluorescence analysis

For immunofluorescence analysis, 3 x 10^4^ VeroE6 cells were seeded in microscopy slide chambers (Falcon; #354108), and incubated overnight. The next day, cells were transduced with either VSV*AG, VSV*ΔG(H3stem), or VSV*AG(HA PR8) at an MOI = 1 in MEM for 12 hours. After washing with PBS, cells were fixed with fixation buffer (3 % paraformaldehyde in PBS) for 20 minutes, quenched (50 mM NH4Cl in PBS) for 10 minutes, and permeabilized in 0.5 % Triton X-100 diluted in PBS for 5 minutes. To block unspecific binding, 2.5 % milk powder (w/v) with 0.1 % Triton X-100 in PBS were added and incubated for 20 minutes on the shaker. Cells were washed once with PBS, and stained with 1:1,000 dilution of ferret α-H1N1 or ferret α-H3N2 serum in 0.1 % Triton X-100 in PBS for 2 hours in the dark. Subsequently, cells were washed again, and secondary antibody dilution (goat α-ferret TxRed-conjugated; 1:1,000 in 0.1 % Triton X-100/PBS) was added for 1.5 hours in the dark. Cells were washed three times with PBS and stained with DAPI (100 ng/mL in water) to visualize cell nuclei. Finally, cells were washed three times with water, mounted in ProLong Diamond Antifade Mountant, covered with a 24×32 mm cover slip, and stored in the dark at 4°C. Pictures were taken at an Axio Observer.Z1 microscope, and analyzed using the ZENblue 3.3 software.

### Confirmation of single-round character of VSV replicons

BHK-G43 cells were seeded in 6-well plates to reach 90 % confluence and treated with 1 nM of mifepristone or left untreated for 6 hours. After incubation period, cells were transduced with a MOI=1 of VSV*ΔG(HA PR8) and incubated at 37°C and 5 % CO_2_ for 24 hours. The next day, 5 μL of supernatant were transferred to a fresh, untreated monolayer of BHK-G43 cells and incubated as described above for another 24 hours. Pictures were taken using a Nikon Eclipse T*i*-S light microscope with software Nis-Elements 4.20.00 LO.

### Animal experiments

All animal experiments were conducted in compliance with the German animal protection laws and authorized by the responsible state authority of Hesse (Regierungspräsidium Darmstadt, protocol F107/1058). Six-to eight-weeks-old female C57BL/6J mice (Jackson Laboratories/Charles River) were immunized i.m. with 1 x 10^6^ ffu of the respective VSV replicons or mixtures thereof, diluted in sterile PBS to a total volume of 30 μl per injection. After four weeks, animals were immunized again using the same dose of the same replicon. Four weeks after the second immunization, animals were anesthetized using intraperitoneal (i.p.) injection of ketamine (100 mg/kg body weight) and xylazine (5 mg/kg body weight) and inoculated i.n. with 6 x 10^2^ or 1.2 x 10^3^ TCID_50_ of PR8 or rSC35M in 30 μL PBS, respectively. During the following 14 days, all animals were monitored daily, and clinical signs of infection were evaluated (Supp. Table 2) ^45^ and body weight was measured. As soon as humane endpoints were reached, mice were euthanized using ketamine (100 mg/kg body weight) and xylazine (10 mg/kg body weight) anesthesia, and bled through cardiac puncture. To investigate underlying humoral immune responses, mice were immunized once or twice as described above and euthanized 28 days after prime or boost immunization by anesthesia and cardiac puncture. Blood was collected in tubes with Z-gel matrix (Sarstedt; Cat#: 41.1378.005), serum was separated by centrifuged for 10 min at 14,000 g and 4°C, and stored at −20°C.

### Total antibody and IgG subtype quantification

Total antibodies specific for influenza virus were measured by immune-peroxidase monolayer assay (IPMA) ^15^. To this end, MDCK cells were seeded into a 96-well plate, incubated at 37°C for 24 hours, and infected with 10^2^ TCID_50_ per well of PR8, or rSC35M the following day. Two days after infection, cells were washed with 1:3 PBS in distilled water, air-dried and heat-fixated at 65°C for 8 hours. Infected MDCK cells were incubated in duplicates by serial two-fold dilutions of mouse sera for 2 hours. After 1 hour, plates were washed three times with 100 μL PBS. Secondary antibodies were added in 100 μL in a 1:750 dilution (Jackson Immunoresearch; total IgG: Cat#: 115-035-003, IgG1: Cat#: 115-035-205, IgG2b: Cat#: 115-035-207, IgG2c: Cat#: 15-035-208, IgG3: Cat#: 115-035-209 and infected cells were visualized as described for virus titration. Antibody titers are expressed as the reciprocal of the highest dilution showing positive staining.

### Neutralizing antibody quantification

For quantification of neutralizing antibodies, 1 x 10^2^ TCID_50_ of either PR8 or rSC35M were incubated with serial two-fold dilutions of serum at room temperature for 20 minutes in a 96-well plate ^15^. Subsequently, 2 x 10^4^ MDCK cells were added. After three days, infected cells were stained as described for virus titration and neutralizing antibody titers were expressed as the reciprocal of the highest dilution showing no infected cells.

### FcγR-IV effector assay

To investigate the capacity of antibodies to activate murine FcγR-IV, the mouse FcγR-IV ADCC Bioassay (Promega) was used. All steps were performed following the manufacturer’s instructions. Briefly, confluent MDCK cells in a 96-well plate were infected with PR8 or rSC35M virus with an MOI = 1 in DMEM and incubated for 18 hours at 37°C (PR8), or 32°C (rSC35M). Sera to be tested were 2-fold diluted (starting dilution 1:10) in a U-bottom plate in 4% assay buffer. After washing the infected cells with PBS, 25 μl of 4% assay buffer and 25 μl of the respective serum dilution was transferred in duplicates. 25 μl of effector cell suspension were added incubated at 37°C for 6 hours. Afterwards, cells and Bio-Glo Reagent were put at ambient temperature for 15 minutes and 75 μl of prepared Bio-Glo Reagent were added to the cells and incubated for 20-30 minutes in the dark at room temperature. 140 μl were then transferred into a black 96-well plate without creating bubbles and luminescence was measured using the PHERAStar FSX luminescence reader.

### ELISpot assays

Spleens of mice immunized with VSV replicons or infected with IAV were taken after euthanasia and single cells were isolated by pressing the organs through a 70 μm cell strainer. Red blood cells were lysed for 1 minute at room temperature using ACK lysis buffer (RECIPE), and remaining cells were resuspended in RPMI and counted. DC2.4 murine dendritic cells were used as stimulator cells and left uninfected or were infected with IAV PR8, rSC35M, or Fluenz Tetra 24 hours prior to the assay. Murine IFNγ and IL-4 single color ELISpot assays were performed using precoated plates (CTL Immunospot) according to the manual. Briefly, 250,000 splenocytes were incubated with 50,000 stimulator cells, or with concanavalin A (ConA) as unspecific stimulus for 40 hours at 37°C. Afterwards, ELISpot plates were washed and treated as described in the manual. Plates were imaged with a Series 6 Universal Analyzer (CTL Immunospot), and analysis was performed using CTL Immunospot Software.

### Statistical analysis

For graphical illustration and statistical analysis, GraphPad Prism 9 was used. Antibody titers were log2-transformed and one- or two-way analysis of variance (ANOVA) with Tukey’s multiple comparison post-test was applied. All groups are compared if not stated otherwise and p-values of <0.05 were considered significant.

### Amino Acid Homology Analysis

Amino acid homology analysis was performed by using ClustalW alignment. Accession numbers of the viral protein sequences used for the alignment are summarized in Supp. Table 4.

## Author Contributions

KW, GZ, CKP – conceptualization; KW, BS, CKP – design of experiments; KW, GZ, MR, SR, YK – data acquisition; KW, CKP – data analysis; KW, GZ, BS, CKP – data interpretation; KW, CKP – writing of the original manuscript; KW, GZ, MR, SR, YK, BS, CKP – editing of the manuscript.

## Acknowledgments

We would like to thank Prof. Dr. Eva Friebertshäuser (Philipps-University Marburg) for kindly providing the rSC35M virus, and her and Dr. Ralf Wagner (Paul-Ehrlich-Institute) for the helpful discussions about this project. We also thank Dr. Katayoun Ayasoufi (Mayo Clinic) for the deep discussions of immunological questions and her helpful comments on the manuscript. We are grateful to Prof. Dr. Karl-Klaus Conzelmann (University of Munich) for providing us with anti-VSV serum. We also thank Dr. Hanna Sediri-Schön (Paul-Ehrlich-Institute) for the help with production of egg-grown virus stocks.

## Data availability

All primary data obtained in this study is included in the supplementary material. Materials generated in this study can be made available to researchers upon request to the corresponding author, but are subject to material-transfer-agreements.

---

**Supplementary Figure 1:**
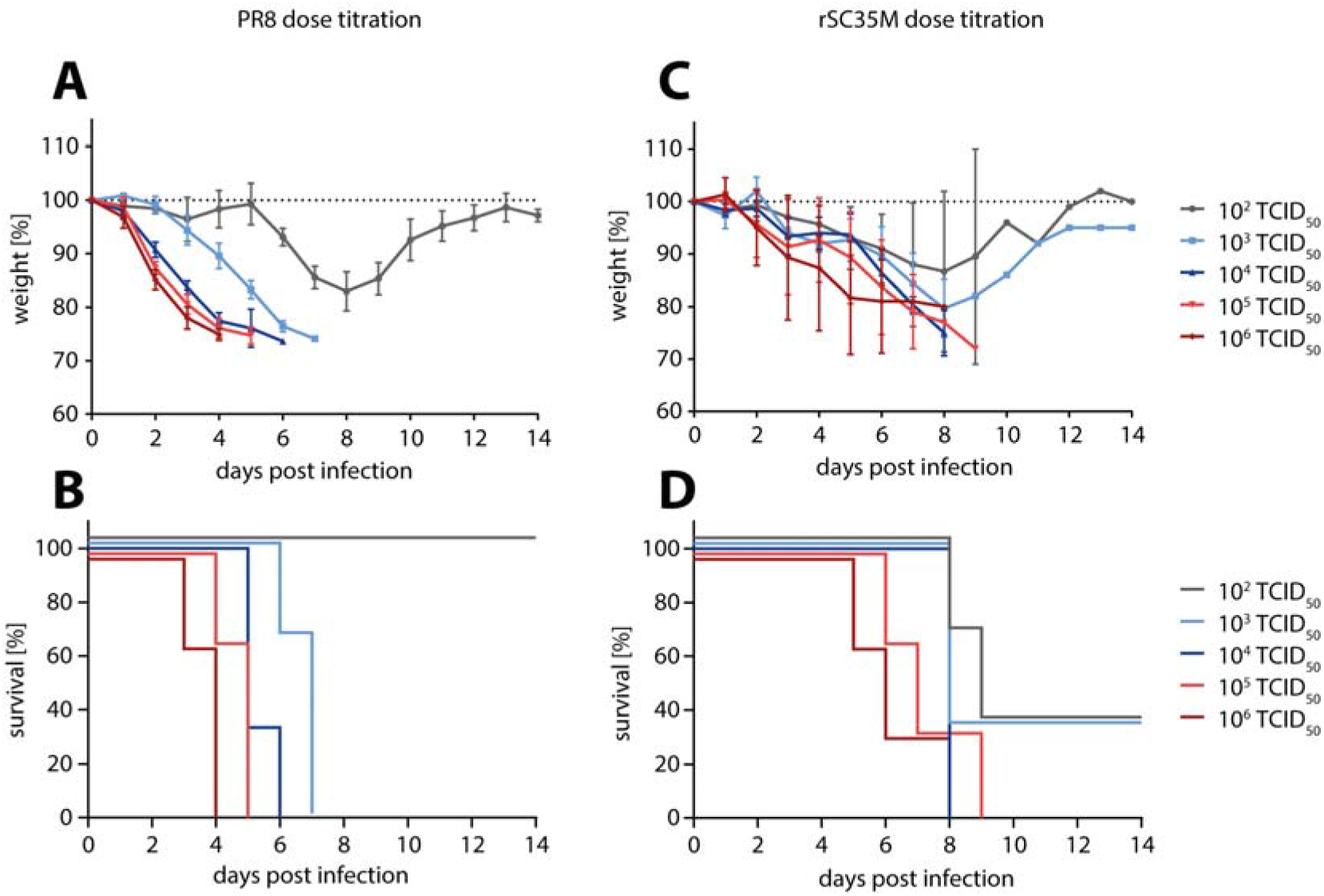
50% lethal dose (LD_50_) titration of PR8 and rSC35M in C57BL/6J mice. Mice were infected with increasing doses of PR8 or rSC35M and monitored daily for 14 days. When reaching 25% weight loss or clinical score of ≥8 (see Supp. Table 2), animals were euthanized. (A) Relative weight loss of PR8 infected mice normalized to start weight on day 0. Dotted line indicates 100%. (B) Survival rate of PR8 infected mice. (C) Relative weight loss of rSC35M infected mice normalized to start weight on day 0. Dotted line indicates 100%. (D) Survival rate of rSC35M infected mice. Data points show mean values and error bars represent standard deviations.

**Supplementary Figure 2:**
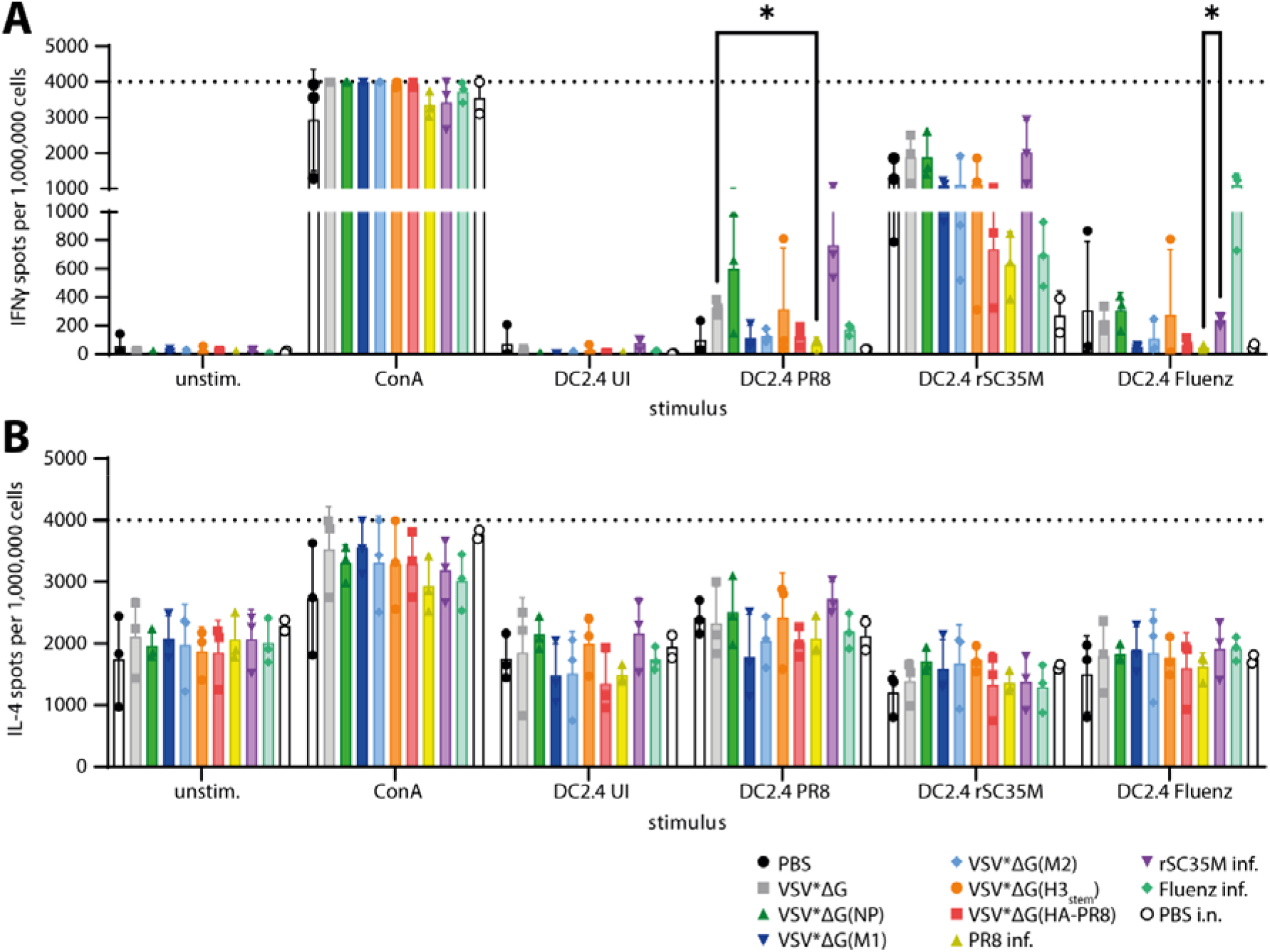
IFNγ- and IL-4-T cell responses of mice vaccinated with VSV replicons or infected with IAV. Mice (n = 3) were immunized with VSV-replicons or PBS intramuscularly in a prime/boost regimen four weeks apart and euthanized one week after boost immunization. Separate groups of mice were intranasally infected with 200 TCID_50_ of PR8, 400 TCID_50_ of rSC35M (equaling 1 LD_50_ for each virus), or with 2,000 TCID_50_ of tissue culture-amplified Fluenz Tetra, and euthanized one week post infection. Splenocytes were isolated and 250,000 cells each were used for (A) murine IFNγ, and (B) murine IL-4 ELISpot assays using a murine dendritc cell line (DC2.4) as stimulator cells either uninfected (UI) or infected with PR8, rSC35M, or Fluenz Tetra for 24 h. Concanavalin A (ConA) was used as unspecific stimulus for positive control, and unstimulated splenocytes (unstim.) served as negative control. Dotted lines indicate upper limit of spot detection (too numerous to count). For statistical analysis, two-way ANOVA was performed followed by Tukey’s multiple comparisons test within each stimulus group. *: p≤0.05.

**Supplementary Figure 3:**
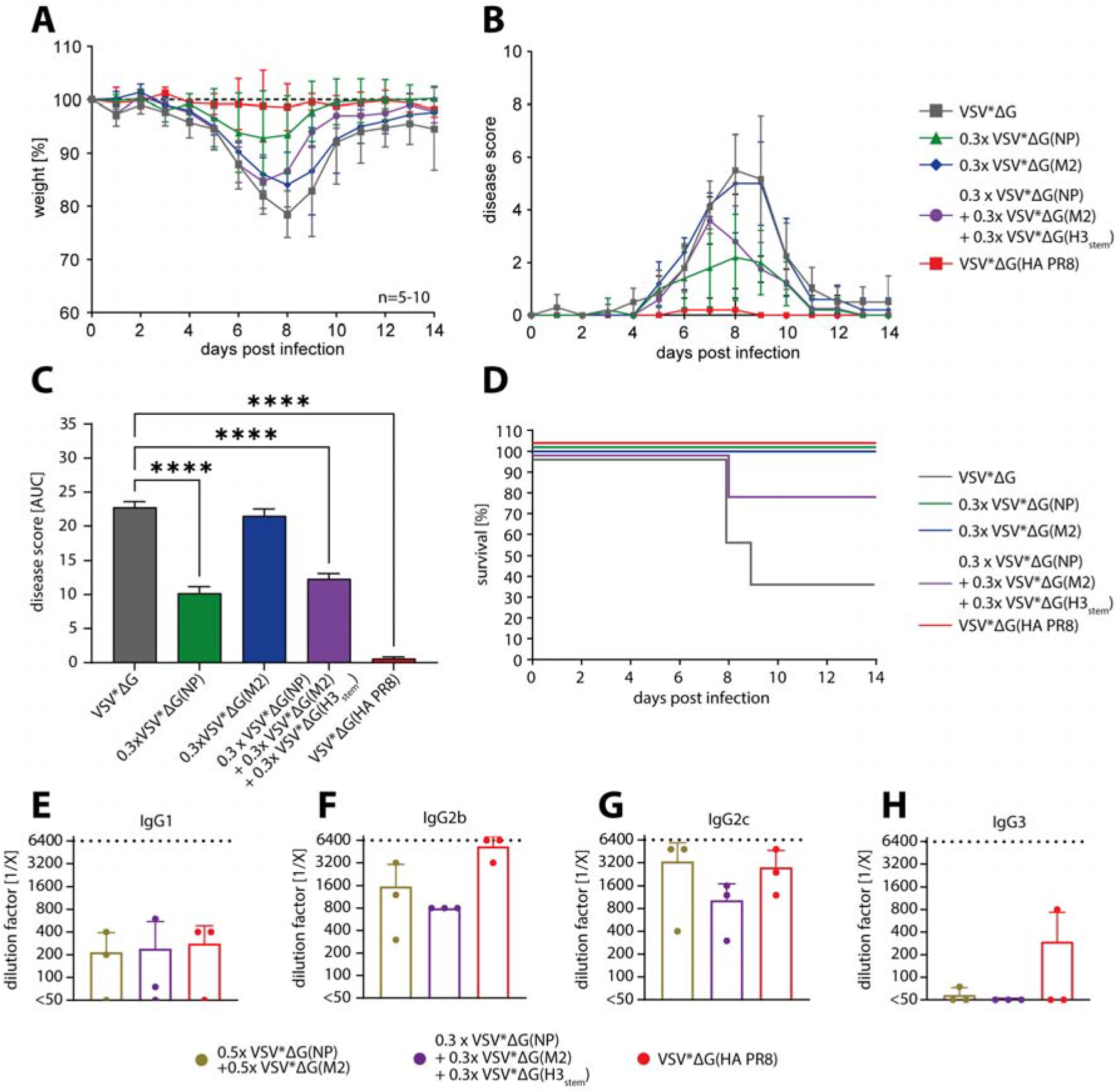
Protection of C57BL/6J mice after prime-boost immunization with VSV replicon triple-cocktail immunizations against PR8. (A) Weight loss of PR8 infected animals during the 14 days after challenge relative to their weight on day 0. Dotted line indicates 100%. Each group consisted of n=5 to 10 mice. VSV*ΔG and VSV*ΔG(HA PR8) groups are the same as in Figure 6. (B) Disease score of PR8 infected mice representing the sum of numerical values for weight loss, behavior, and appearance. For details, see Supp. Table 2. Data points in (A) and (B) represent means of the respective groups, and error bars indicate standard deviations. (C) Area under curve (AUC) analysis of disease scores. Height of bars represent the mean and error bars are standard error of the mean. For statistical analysis, one-way analysis of variance with Tukey’s multiple comparison post-hoc-test using the VSV*ΔG group as a reference was performed; ****p<0.0001. (D) Survival of mice infected with PR8. Surviving mice were euthanized at the end of the study. (E-H) PR8-reactive IgG subclass analysis of double-cocktail (gold), triple-cocktail (purple), and VSV*ΔG(HA PR8) immune sera (red) 28 days after boost vaccination. Last wells showing positive staining are expressed as the reciprocal of the serum dilution factor. Bars represent the mean, and error bars are standard deviations. The dotted line represents the upper limit of detection.

**Supplementary Figure 4:**
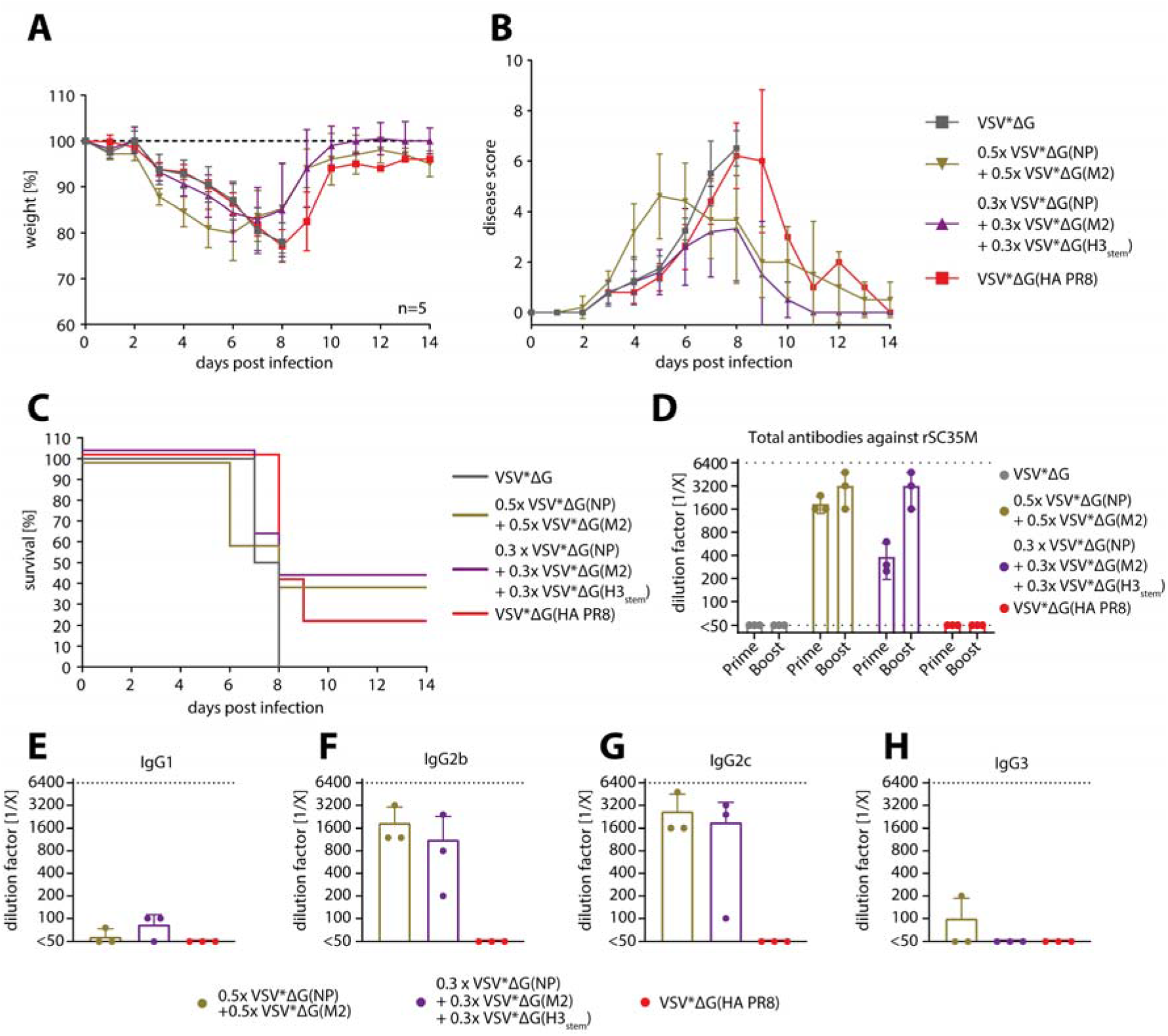
Protection of C57BL/6J mice after prime-boost immunization with VSV replicon double- and triple-cocktail immunizations against rSC35M. (A) Weight loss of rSC35M infected animals during the 14 days after challenge relative to their weight on day 0. Dotted line indicates 100%. Each group consisted of n=5 mice. (B) Disease score of rSC35M infected mice representing the sum of numerical values for weight loss, behavior, and appearance. For details, see Supp. Table 2. Data points in (A) and (B) represent means of the respective groups, and error bars indicate standard deviations. (C) Survival of mice infected with rSC35M. Surviving mice were euthanized at the end of the study. (D-H) Antibody titrations of immune sera 28 days after boost vaccination against rSC35M. Last wells showing positive staining are expressed as the reciprocal of the serum dilution factor. Dotted lines represent the lower and upper limits of detection, respectively. Height of bars indicate mean of groups and error bars are standard deviations. (D) Total IgG titration. (E-H) IgG subclass analysis of double-cocktail (gold), triple-cocktail (purple), and VSV*ΔG(HA PR8) (red).

